# Neoadjuvant anti-4-1BB confers protection against spontaneous metastasis through low-affinity intratumor CD8^+^ T cells in triple-negative breast cancer

**DOI:** 10.1101/2025.01.29.635356

**Authors:** Bryan Jian Wei Lim, Mingyong Liu, Liuyang Wang, Sharleen Li Ying Kong, Tao Yin, Chengsong Yan, Kun Xiang, Chengjie Cao, Haiyang Wu, Ariana Mihai, Felicia Pei-Ling Tay, Ergang Wang, Qizhou Jiang, Zhehao Ma, Lianmei Tan, Rui Ning Chia, Diyuan Qin, Christopher C. Pan, Xiao-Fan Wang, Qi-Jing Li

**Affiliations:** Department of Integrative Immunobiology, Duke University School of Medicine, Durham, NC, USA; Department of Pharmacology and Cancer Biology, Duke University School of Medicine, Durham, NC, USA; Institute of Molecular and Cell Biology, Agency for Science, Technology and Research (A*STAR), Singapore, Singapore; Department of Molecular Genetics and Microbiology, Duke University School of Medicine, Durham, NC, USA; TCRCure Biopharma, Durham, NC, USA; Duke Kunshan University, Kunshan, China; Duke-NUS Medical School, Singapore, Singapore; Singapore Immunology Network, Agency for Science, Technology and Research (A*STAR), Singapore, Singapore

**Author notes:** Corresponding authors: Qi-Jing Li,; Xiao-Fan Wang.

**Keywords:** Neoadjuvant, immunotherapy, agonistic, 4-1BB, exhaustion, intermediate, transitory, metastasis, systemic, affinity

## Abstract

Neoadjuvant immunotherapy seeks to harness the primary tumor as a source of relevant tumor antigens to enhance systemic anti-tumor immunity through improved immunological surveillance. Despite having revolutionized the treatment of patients with high-risk early-stage triple-negative breast cancer (TNBC), a significant portion of patients remain unresponsive and succumb to metastatic recurrence post-treatment. Here, we found that optimally scheduled neoadjuvant administration of anti-4-1BB monotherapy was able to counteract metastases and prolong survival following surgical resection. Phenotypic and transcriptional profiling revealed enhanced 4-1BB expression on tumor-infiltrating intermediate (T_int_), relative to progenitor (T_prog_) and terminally exhausted (T_term_) T cells. Furthermore, T_int_ was enriched in low-affinity T cells. Treatment with anti-4-1BB drove clonal expansion of T_int_, with reduced expression of tissue-retention marker CD103 in T_prog_. This was accompanied by increased TCR clonotype sharing between paired tumors and pre-metastatic lungs. Further interrogation of sorted intratumor T cells confirmed enhanced T cell egress into circulation following anti-4-1BB treatment. In addition, gene signature extracted from anti-4-1BB treated T_int_ was consistently associated with improved clinical outcomes in BRCA patients. Combinatorial neoadjuvant anti-4-1BB and ablation of tumor-derived CXCL16 resulted in enhanced therapeutic effect. These findings illustrate the intratumor changes underpinning the efficacy of neoadjuvant anti-4-1BB, highlighting the reciprocity between local tissue-retention and distant immunologic fortification, suggesting treatment can reverse the siphoning of intratumor T cells to primary tumor, enabling redistribution to distant tissues and subsequent protection against metastases.

## INTRODUCTION

Triple-Negative Breast Cancer (TNBC) is an aggressive subtype that comprises approximately 15 to 20% of all breast cancer cases^1,2^. It is a biologically heterogenous disease, clinically characterized by absence of estrogen receptor (ER) and progesterone receptor (PR), accompanied by lack of amplification of human epidermal growth factor receptor 2 (HER2)^3–5^. Without access to targeted therapies, patients are presented with a paucity of treatment options, thereby limiting them to the use of only chemotherapy which remains the standard therapeutic approach for TNBC patients^3,5^. However, a significant proportion of patients harbor residual disease and experience metastatic recurrence^6–8^. Because of the high risk of metastasis and recurrence post-treatment, there has been an unmet clinical need to address these challenges^9–11^.

Neoadjuvant therapy is treatment given prior to surgery to downstage disease and improve surgical prognosis^12^. With the paradigm shifting results from KEYNOTE-522, the clinical management of high-risk early-stage TNBC has progressed to include anti-PD-1 in the neoadjuvant setting in combination with chemotherapy^13–16^. It is now recognized that high levels of tumor infiltrating lymphocytes (TILs) are not only associated with improved prognosis^17^, but also predictive of responsiveness to immunotherapy^18,19^. Neoadjuvant immune checkpoint blockade (ICB) posits that reinvigorated cytotoxic TILs can egress and eliminate micrometastatic tumors that previously disseminated^20–22^. However, many patients do not show an adequate response to affect their long-term survival^11,13–15^. In fact, approximately 50% of TNBC patients who do not achieve pathological complete response (pCR) will experience recurrence^11^. Furthermore, most cancer therapy studies conducted in animals to date have focused locally on the primary tumor, although metastatic recurrence remains the most common cause of death for patients^23–25^.

Amongst the TILs, CD8^+^ T cells play a critical role in mediating anti-tumor cytotoxic functions targeted at cancer cells^26^. However, chronic antigen stimulation within the tumor microenvironment (TME) skews T cells towards T cell exhaustion, a state characterized by gradual loss of effector function and upregulation of co-inhibitory receptors^27,28^. Multiple studies have proposed a model in which exhausted T cells (T_ex_) differentiate through a linear ontogeny, progressing from upstream progenitor (T_prog_) to downstream intermediate (T_int_), and ultimately terminal (T_term_) subsets, each with distinct roles and characteristics in anti-tumor immunity^29–33^. Furthermore, recent reports demonstrated that some infiltrating T_ex_ cells taking up resident markers^34–37^. The presence of such T cells correlates positively with prognosis in human breast cancer^34,38^. It is now known that intratumor CD8^+^ T cells are heterogenous in composition, with only a limited pool that responding to ICB^26,30,34,39,40^. Importantly, T_ex_ have been shown to be early responders to ICB, egressing following combinatorial treatment with anti-TGFβ through downregulation of tissue-retention marker CD103 and correlating with favorable prognosis^41,42^. Indeed, CD8^+^ T cells are capable of functioning as effective sentinels beyond the primary tumor, combating tumor cells at multiple points along the metastatic cascade^43,44^.

The effectiveness of immunotherapy is dependent on successful induction of a systemic immune response^45^, with the composition of T cell repertoire of TILs contributing to prognosis for survival^46^. The tumor-reactive T cell pool is highly diverse and comprised of TCR of varying affinities^47–49^. Due to the process of negative selection in the thymus, majority of T cells targeting tumor-associated antigens (TAAs) express lower affinity TCR^50^. Despite low-affinity TCRs being associated with memory generation and superior persistence^51,52^, they are vulnerable to suppressive cues within the TME^53^. Therefore, it is imperative to identify candidates for immunotherapy to target in the neoadjuvant setting with the intention of promoting egress of low-affinity T cells prior to surgical resection, thereby bolstering immunosurveillance against metastatic recurrence.

In this study, we identified 4-1BB (CD137 or *Tnfrsf9*) as a suitable candidate for neoadjuvant immunotherapy. 4-1BB is a potent costimulatory molecule, expressed on T cells only after activation^54,55^. In human cancers, 4-1BB can serve as a biomarker for tumor-specific CD8^+^ T cells, capable of enriching for full repertoire of antigen-reactive pool without prior knowledge of epitope specificities^56,57^. Over the years, the use of agonistic antibodies to target 4-1BB has gained considerable traction in the translation space^58,59^. However, implementation of agonistic antibodies targeting human 4-1BB in the clinic required managing trade-offs between liver toxicity and efficacy that ultimately led to cessation of trials^59^. Nevertheless, ex vivo studies have shown that anti-4-1BB could enhance the Notably, anti-4-1BB in the neoadjuvant setting has never been tested in human TNBC clinical trials. Moreover, whether anti-4-1BB can drive egress of T cells to augment protection in the metastatic phase of disease remains undetermined.

Herein, we demonstrate that neoadjuvant anti-4-1BB is protective against spontaneous metastasis. 4-1BB is enriched in T_int_ and more broadly, low-affinity T cells. Anti-4-1BB can promote clonal expansion of T cells within the primary tumor and more specifically T_int_. Finally, we found that anti-4-1BB promotes T cell egress of expanded T_ex_ subsets from the primary tumor thereby increasing clonotype sharing with lungs and to a lesser extent, the spleen but not the contralateral fatpad. These findings suggest that anti-4-1BB is a potential therapeutic target in the neoadjuvant setting to counter metastatic recurrence in TNBC.

## RESULTS

### Neoadjuvant anti-4-1BB monotherapy prolongs survival against spontaneous metastases

Surgery remains the major treatment modality in the clinic for early-stage TNBC^60,61^. Mouse studies have demonstrated that the presence of the primary tumor acts as an immune suppressor, with its resection leading to increased immune competence despite the presence of metastatic disease^62,63^. Using the spontaneously metastatic 4T1 model, which is highly resistant ICB^64,65^, we first determined the validity of this model to recapitulate the lack of response to anti-PD-1. Due to the proximity of the inguinal lymph node and mammary tumor growth site, we performed both lymphadenectomy and primary tumor removal on day of surgical resection (Figure S1A). In agreement with previously published results^64,65^, we found that anti-PD did not affect primary tumor growth (Figure S1B, C). More importantly, we also demonstrated the lack of efficacy of anti-PD-1 in countering metastatic disease post-surgical resection (Figure S1D, E). Previous studies have indicated the importance of timing between neoadjuvant immunotherapy and surgery^66^. As such, we repeated the experiment with a 4-day window between the start of treatment to surgery (Figure S1F). However, this neither affected metastasis nor primary tumor growth (Figure S1G-J), unlike what was previously found in 4T1.2^67^, demonstrating the relatively aggressive nature of parental 4T1 line.

Besides checkpoint molecules, TILs express another array of molecules termed co-stimulatory molecules which can be targeted with agonistic antibodies^68–70^. As such, we analyzed our previously published scRNA-seq of sorted tumor-infiltrating TCRβ^+^CD44^+^CD62L^-^CD69^-^CD103^-^ T_EFF/EM_ (effector/effector memory) from 4T1 primary tumors^71^. We performed uniform manifold approximation and projection (UMAP) for dimensionality reduction (Figure S2A, B). Focusing only on CD8^+^ T cell enriched subsets, we obtained 7 clusters after removing CD4^+^ T cells (Figure 1A). Of all the costimulatory molecules analyzed, 4-1BB was the only candidate that was strongly expressed in cluster 5 which was enriched for *Pdcd1* (PD-1) and *Cxcr6* (CXCR6) (Figure 1B). Importantly, our group previously demonstrated the anti-metastatic potential of CXCR6^+^PD-1^+^CD8^+^ T cells^71^. As with anti-PD-1, we determined the efficacy of neoadjuvant anti-4-1BB as monotherapy, using the same experimental setup (Figure S2C). Anti-4-1BB did not cause significant difference in terms of primary tumor growth (Figure S2D). Importantly, mice given anti-4-1BB significantly prolonged survival relative to isotype-treated group (Figure S2E), indicating this protection was not attributable to differences in size of primary tumors. This benefit is supported by another study which employed the TNBC E0771 model and showed a similar survival benefit of anti-4-1BB relative to isotype-treated group, though primary tumor volume was not reported^67^. Enumeration of metastatic 4T1 tumor nodules on lung surface at individual endpoints also showed reduced lung metastatic burden in anti-4-1BB treated group (Figure S2F). Given that 4-1BB signaling displays delayed kinetics^72^, we extended the window between the start of anti-4-1BB treatment to surgery to 6 days (Figure 1C). Once again, we noted no significant difference in primary tumor growth (Figure 1D). However, through extension of the neoadjuvant window, 50% of animals in anti-4-1BB treated group were able to achieve long-term survivors (Figure 1E) that showed no signs of metastases by the end of study (Figure 1F).

**Figure 1.**
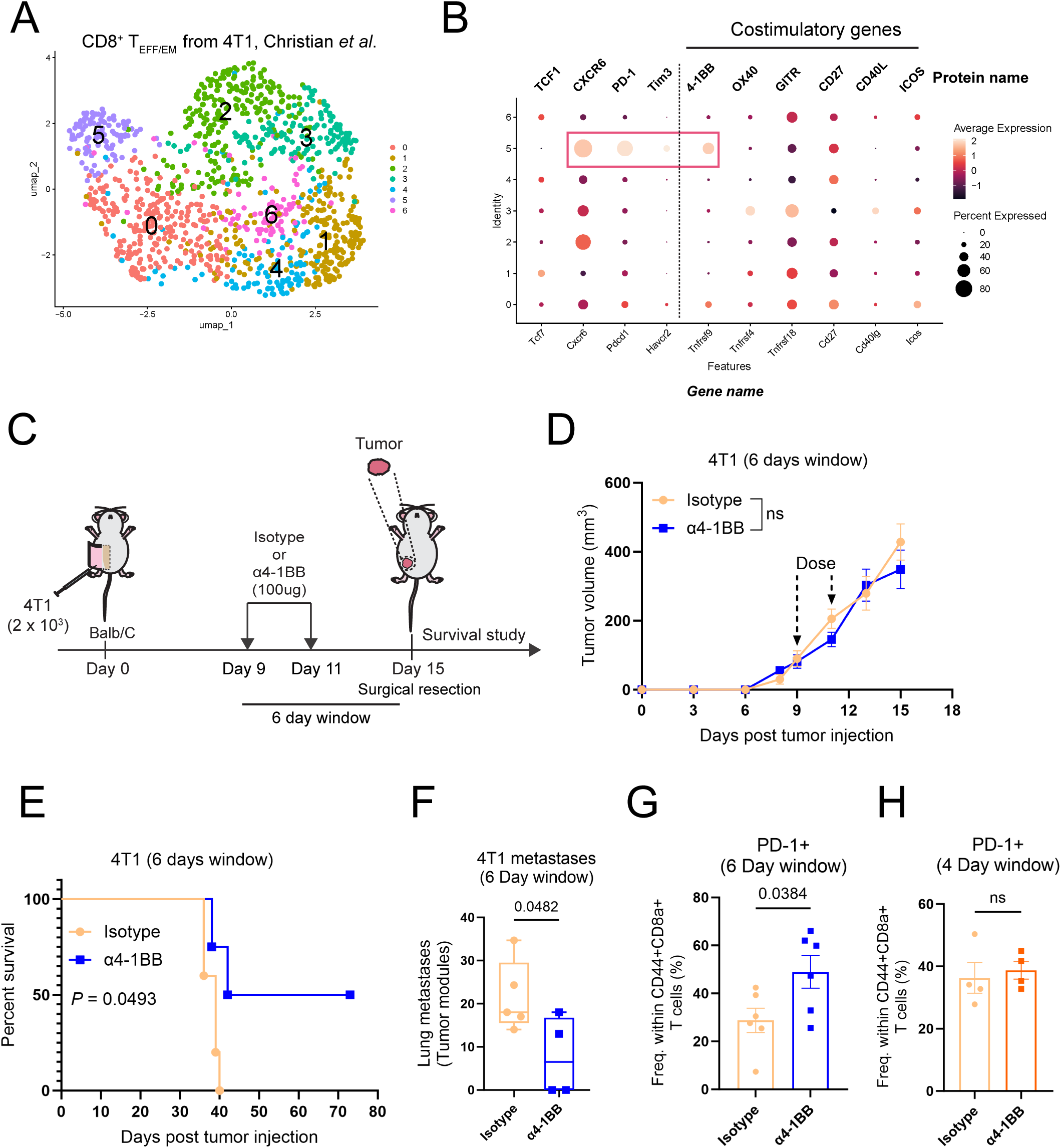
Neoadjuvant anti-4-1BB prolongs survival in spontaneous metastasis murine model following expansion of PD-1^+^CD8^+^ TILs. **(A)** UMAP of CD8^+^ T_EFF/EM_ only from 4T1 tumors conducted by Christian. *et al*. **(B)** Dotplot showing selective co-expression of 4-1BB (*Tnfrsf9*), and no other co-stimulatory genes, with common exhaustion markers CXCR6 (*Cxcr6*), TCF1 (*Tcf7*), PD-1 (*Pdcd1*) and TIM3 (*Havcr2*). **(C-G)** 6**-**day window experimental timeline for orthotopic 4T1 metastasis model. **(C)** Experimental schematic. **(D)** Primary tumor growth. **(E)** Kaplan-Meier analysis. **(F)** Lung metastatic nodule counts at endpoints. **(G)** Frequency of PD-1^+^CD8^+^ T cells in tumor. **(H)** Frequency of PD-1^+^CD8^+^ T cells in tumor under 4-day window. (in D, Isotype, n = 7; α4-1BB, n = 7; in E-F, Isotype, n = 5; α4-1BB, n = 4; in G, Isotype, n = 6; α4-1BB, n = 6; in H, Isotype, n = 4; α4-1BB, n = 4.) Data are presented as mean ± SEM. *P* values were determined using two-way repeated measures ANOVA (D), log-rank (Mantel-Cox) test (E), two-tailed unpaired Student’s *t*-test (F-H); ns, not significant.

Late diagnosis presents a significant challenge in the clinic, often requiring adjustments to treatment regimen. By delaying commencement of treatment (Figure S3A), we sought to determine if delaying initial dosing would negatively impact the therapeutic effect of neoadjuvant anit-4-1BB. Consistent with our earlier experiments, we saw that primary tumor remains uncontrolled (Figure S3B, C). However, neoadjuvant anti-4-1BB treated groups still trended towards a significant survival benefit, though it was accompanied by diminished metastatic control as indicated by similar lung metastatic burden between isotype and anti-4-1BB treated groups (Figure S3D, E). To determine if T cells contribute to the survival benefit mediated by anti-4-1BB, we employed immune-deficient *Rag2KO* Balb/C mice. We noted no changes in primary tumor burden and more importantly, no survival benefits nor metastatic protection across repeated experiments (Figure S3F-N), confirming the importance of T cells in protecting against metastases. CD8^+^ T cells have been reported to be the primary responders and effectors following anti-4-1BB treatment^73–75^. Importantly, when we analyzed tumor-infiltrating PD-1^+^CD44^+^CD8^+^ T cells, we noted significant increase only in the 6-day as opposed to the 4-day window (Figure 1G, H). Taking this, together with the CD8^+^ T cell dependent metastatic protection, it alludes to the importance of expansion of PD-1^+^CD8^+^ T cells in the tumor for subsequent protection against metastases. In summary, we have demonstrated the capability of anti-4-1BB, and not anti-PD-1, to function as a monotherapy in the neoadjuvant setting to control spontaneous metastases, likely through CD8^+^ T cells.

### Expression of 4-1BB is enriched in T_int_ within tumor

To determine the origins of T cells responding to neoadjuvant anti-4-1BB, we used flow cytometry to assess the expression level of 4-1BB in CD8^+^ T cells from the tumor and dLN. Notably, 4-1BB was enriched only in tumor and not dLN across multiple TNBC mouse models (Figure 2A, B). Likewise, the exhaustion molecule PD-1 displayed a similar expression pattern (Figure 2C, D). Indeed, we confirmed higher levels of 4-1BB in PD-1^+^CD8^+^ T cells as opposed to PD-1^-^CD8^+^ T cells (Figure 2E). TILs undergo chronic antigen stimulation, driving them towards exhaustion^27^. Given that 4-1BB is upregulated only after T cell activation^54^, its exclusive presence in the tumor, and parallel expression with exhaustion marker PD-1 potentially reflects the availability of antigens and represent a consequence of chronic activation. TCF1 and TIM3 are canonical markers used to identify distinct subsets of T_ex_ in both chronic viral infection and cancer models in mice, namely the T_prog_ and T_term_ respectively^29,40,76^. Importantly, this annotation translates to human cancer^77–79^. It was established that TCF1 is required to maintain stemness of the non-cytotoxic T_prog_^80^, and attenuation of TCF1 leads to loss of stemness and subsequent gain in cytotoxic effector functions as T_prog_ transitions into T^81,82^.

**Figure 2.**
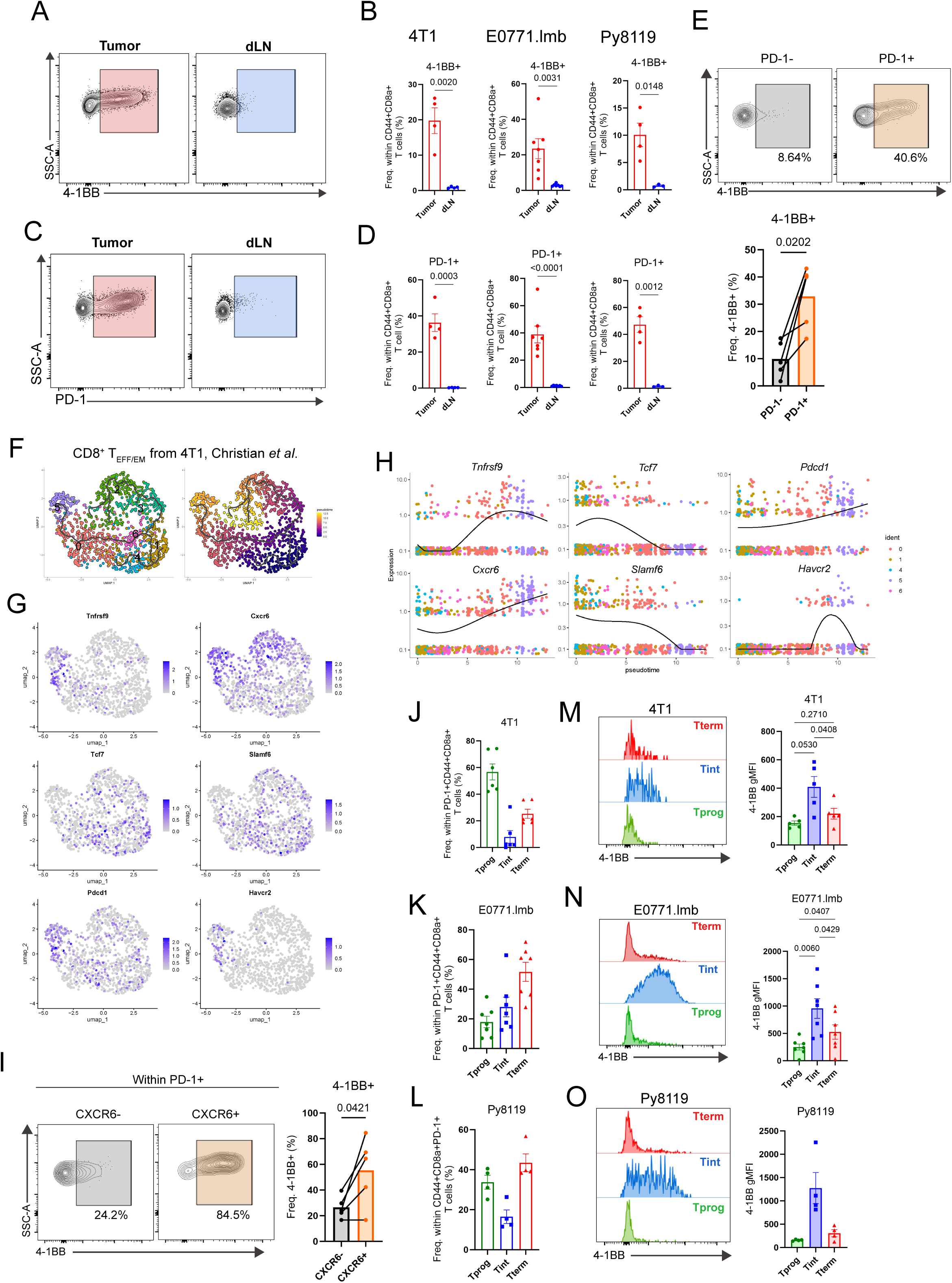
4-1BB is enriched in intermediate exhausted T cells (T_int_) of tumor in multiple TNBC mouse models. (A-B) 4-1BB is selectively expressed in T cells from tumor and not dLN. **(A)** Representative flow plots. **(B)** Frequency of 4-1BB in tumor and dLN CD8^+^ T cells across 4T1, E0771.lmb, Py8119 tumor models at Day 15. **(C-D)** PD-1 is selectively expressed in T cells from tumor and not dLN. **(C)** Representative flow plots. **(D)** Frequency of PD-1 in tumor and dLN CD8 T cells across 4T1, E0771.lmb, Py8119 tumor models. **(E)** Expression of 4-1BB in PD-1^-^ and PD-1^+^ CD44^+^CD8^+^ T cells from 4T1 tumors, representative flow plot (top), quantification (bottom) (n = 4). **(F)** UMAP (left) of CD8^+^ T_EFF/EM_ only from 4T1 tumors conducted by Christian. *et al* with accompanying pseudotime trajectory (right) showing progression of CD8^+^ T cell clusters (from 4T1 tumor) in development. TCF1^+^ subset (cluster 1) was used as the starting point in trajectory analyses. **(G)** Feature plots showing expression of 4-1BB (*Tnfrsf9*), CXCR6 (*Cxcr6*), TCF1 (*Tcf7*), LY108 (*Slamf6*), PD-1 (*Pdcd1*) and TIM3 (*Havcr2*). **(H)** Pattern of *Tcf7*, *Slamf6*, *Tnfrsf9*, *Cxcr6*, *Pdcd1* and *Havcr2* gene expression in clusters 0, 1, 4, 5 and 6 of CD8^+^ T_EFF/EM_ only analysis was projected onto pseudotime space to visualize divergent and parallel gene expressions. **(I)** Expression of 4-1BB in CXCR6^-^PD-1^+^ and CXCR6^+^PD-1^+^ CD44^+^CD8^+^ T cells from 4T1 tumors, representative flow plot (left), quantification (right) (n = 4). **(J-L)** Frequency of T_prog_, T_int_ and T_term_ in tumors of 4T1 (J), E0771.lmb (K), Py8119 (L) at Day 15. **(M-O)** Representative flow plots showing higher 4-1BB expression in T_int_ relative to T_prog_ and T_term_ (left), and gMFI of 4-1BB (right) from models 4T1 (M), E0771.lmb (N), Py8119 (O) at Day 15. (in B, D, J-O, n = 3-5). Data are presented as mean ± SEM. *P* values were determined using two-tailed unpaired Student’s *t*-test (B, D), two-tailed paired Student’s *t*-test (E, I), one-way repeated measures ANOVA followed by post-hoc Tukey’s multiple comparisons test (M-N).

To clearly illustrate the changes in patterns of gene expression during transition of T_ex_ development, we projected cells onto a pseudotime trajectory – setting cluster 1, marked by the highest level of *Tcf7* (TCF1) and *Slamf6* (LY108) expression, indicative of T_prog_, as the starting point (Figure 2F, G). We noted a branch point when cells transitioned between clusters 1 to 3 and 6 with cluster 5 being at one of the endpoints (Figure 2F). Based on *Havcr2* (TIM3) expression being enriched in cluster 5 (Figure 2G), we used only clusters 0, 1, 4, 5 and 6 for plotting gene expression in pseudotime (Figure 2H). Pseudotime analyses further highlight the divergent expression profile of 4-1BB compared to TCF1 and LY108. Conversely, it demonstrates that the *Tnfrsf9* (4-1BB) expression pattern parallels those of PD-1, TIM3 and CXCR6 (Figure 2I). Co-stimulation within the TME was recently shown to be required for effector differentiation of T_prog_ to downstream subsets following its entry from the tumor dLN^83^ (Figure S4A). In support of our findings, recent reports have demonstrated that expression of CXCR6 is repressed by TCF1, and that TCF1 downregulation results in subsequent increase of CXCR6 expression in effector subsets downstream of T_prog_^79,84^. Furthermore, we confirmed that that 4-1BB was more strongly expressed in the CXCR6^+^, and not CXCR6^-^, subsets within the PD-1^+^CD44^+^CD8^+^ T cells at the protein level (Figure 2I).

Recent studies have identified T_int_, denoted by CX3CR1 – a marker of differentiation in antigen experienced T cells – that is functionally distinct from TIM3^+^ T_term_^31,32,85–87^. Given the co-expression patterns of the co-stimulatory molecule 4-1BB with other exhaustion markers described above, we investigated the levels and specificity of 4-1BB expression across the T_prog_ (TCF1^+^/LY108^+^TIM3^-^), T_int_ (TCF1^-^/LY108^-^CX3CR1^+^TIM3^+^) and T_term_ (TCF1^-^/LY108^-^CX3CR1^-^TIM3^+^) subsets using the gating strategy described (Figure S4B). When TIM3 is not present in the panel, we used only TCF1/LY108 and CX3CR1 to delineate the three populations as previously published^31,32,88^. We noted the presence of the three subsets within the PD-1^+^ T_ex_ across multiple TNBC models when tumors were harvested at Day 15 (Figure 2J-L) and Day 21 (Figure S4C-F). Notably, 4-1BB exhibited the highest level of expression in T_int_ over T_prog_ and T_term_ in all models tested (Figure 2M-O). Furthermore, this feature of 4-1BB enrichment in T_int_ was further highlighted when tumors were permitted to grow further (Figure S4G, H). This data suggested that T_int_ was likely the major responders to anti-4-1BB therapy.

### Low-affinity T cells are skewed towards T_int_ and enriched for 4-1BB

TCR affinity has been shown to correlate with the developmental trajectories of T cells in the context of chronic infection models^89^. Furthermore, it has been shown to be a key determinant governing functional T cell response within the TME^90,91^. Low-affinity T cells, which recognize antigens with lower binding strength, play a unique role in the immune response^92^. Despite their lower affinity, these T cells can be highly functional and contribute significantly to immune surveillance and treatment response^51,52,93^. With that in mind, we sought to determine if 4-1BB^+^ T cells were enriched in low or high-affinity T cells. We overexpressed OVA (SIINFEKL) in E0771 and implanted them orthotopically to the fatpad of mice. Using the H2-K^b^-SIINFEKL tetramer, we detected antigen-specific CD8^+^ T cells from OVA expressing tumors. Using tetramer staining intensity as a proxy for TCR affinity^92,94^, we subdivided PD-1^+^CD44^+^CD8^+^ T cells into four groups – tetramer negative, low, mid and high (Figure 3A). We found that 4-1BB was more prominently expressed in tetramer^low/mid^ cells as opposed to tetramer^negative/high^ cells (Figure 3B, C). These results suggested that low-affinity antigen specific T cells are likely responders in neoadjuvant anti-4-1BB.

**Figure 3.**
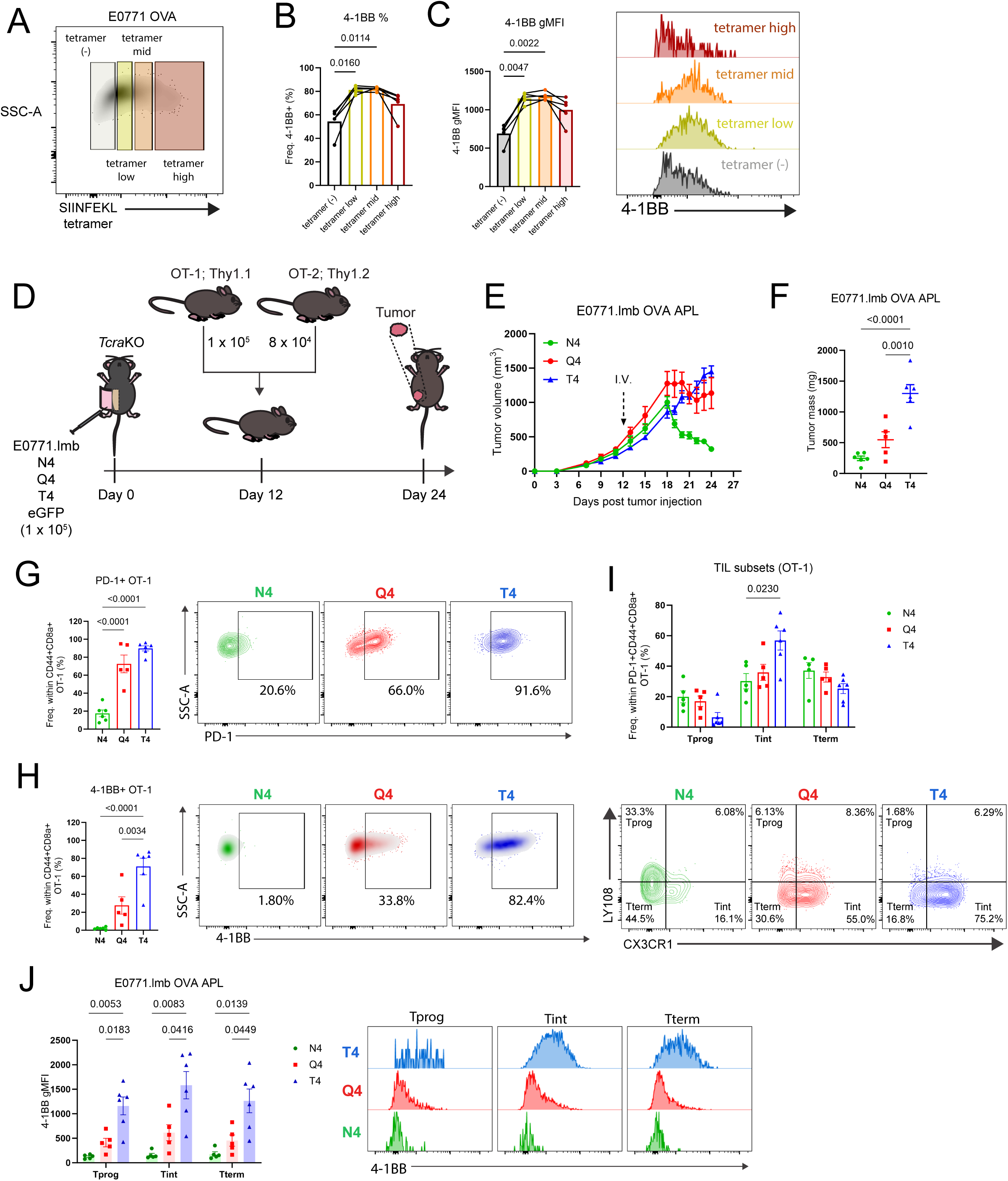
T_int_ and 4-1BB are enriched in low-affinity T cells. (A-C) Low SIINFEKL tetramer levels correlate with higher levels 4-1BB. **(A)** Representative level of SIINFEKL tetramer staining in CD44^+^CD8^+^ T cells infiltrating E0771 OVA tumors. **(B)** Correlation of 4-1BB percentage and **(C)** gMFI level by tetramer intensity in E0771 OVA model, quantification (left), representative flow plots (right). **(D-J)** T_int_ is enriched in low-affinity T cells in E0771.lmb. **(D)** Experimental schematic. **(E)** Tumor volume of E0771.lmb expressing OVA APL following adoptive transfer of OT-1 and OT-2. **(F)** Tumor mass at endpoint of study. **(G)** Frequency (left) and representative FACs plots (right) of PD-1^+^ OT-1. **(H)** Frequency (left) and representative FACs plots (right) of 4-1BB^+^ OT-1. **(I)** Frequency of T_prog_, Tint and T_term_ in PD-1^+^ compartment of OT-1 (top), representative flow plots (bottom). **(J)** gMFI of 4-1BB in T_prog_, T_int_ and T_term_ (left). Representative flow plots showing 4-1BB expression (right) (in E-J, n = 5-6). Data are presented as mean ± SEM. *P* values were determined using one-way repeated measures ANOVA followed by post-hoc Tukey’s multiple comparisons test (B, C), one-way ANOVA followed by post-hoc Tukey’s multiple comparisons test (F-H), two-way ANOVA followed by post-hoc Tukey’s multiple comparisons test (I, J); ns, not significant.

Given that the tetramer binding threshold is higher than that of T cell activation, tetramer staining might potentially miss very weak clones in an open repertoire^95^. Considering how majority of T cells specific for TAA express low-affinity TCRs^47^, we sought to utilize a more sophisticated system to validate the hypothesis that 4-1BB is enriched in low-affinity T cells. To do so, we adopted the SIINFEKL variants or altered peptide ligands (APL) system where the sensitivity of OT-1 for the APLs expressed in EC_50_ of APL/N4 was previously measured (SIIQFEKL EC_50_ = 18.3, SIITFEKL EC_50_ = 70.7, SIIVFEKL EC_50_ = 680)^96^.

To test our hypothesis in vivo, we introduced the OVA APL system by lentiviral transduction into TNBC tumor cells for stable overexpression of the OVA_257-265_ variants, including the constant OVA_329-337_ (AAHAEINEA) (Figure S5A-C). We sought to test the hypothesis that 4-1BB was enriched in low-affinity T cells. To do so, we compared E0771.lmb-N4 (high-affinity), Q4 (moderate-affinity) and T4 (low-affinity) expressing tumors. Tumors were first implanted into *TcrαKO* mice and allowed to grow for 12 days prior to adoptive transfer of T cells (Figure 3E). CD4^+^ T cells have been shown to be important in development of proper effector response in CD8^+^ T cells in chronic LCMV and cancer studies^32,97^. As such, we transferred both OT-1 and OT-2 cells. Adoptively transferred T cells were able to control N4 and Q4 but not T4-expressing tumors, which is attributable to the difference in affinity for OT-1 TCR (Figure 3F). We noticed a 4-fold increase in frequency of PD-1^+^OT-1 cells within the CD44^+^CD8^+^ compartment in Q4 and T4 tumors as opposed to N4 tumors (Figure 3G). Importantly, 4-1BB expression pattern showed a similar inverse relationship with TCR affinity (Figure 3H). When comparing the T_ex_ subsets within the PD-1^+^OT-1 TILs, we found that T_int_ was significantly higher in T4 tumors, taking over the space occupied by T_prog_ (Figure 3I). Most strikingly, 4-1BB expression was significantly enriched across all OT-1 TIL subsets in T4 relative to N4 and Q4 tumors (Figure 3J). To determine if 4-1BB enrichment in low-affinity T cells is not model-specific, we repeated this experiment comparing Py8119-N4 and T4 expressing tumors (Figure S5D). In agreement with our findings on the E0771.lmb model, we recapitulated the findings of low-affinity TCR favoring T_int_ formation and 4-1BB expression (Figure S5E-I). Taken together, the data suggested that moderate and low-affinity T cells are skewed towards T_int_ while broadly displaying the greatest potential for targeting by anti-4-1BB.

### Anti-4-1BB drives systemic dissemination of T_ex_ to distant pre-metastatic tissues

Next, we sought to determine the mechanism in which neoadjuvant anti-4-1BB mediates protection against metastasis. Continuous T cell trafficking of T cells between the tumor and lymphoid tissues have been reported^98,99^. However, chronic antigen stimulation has been shown to confine T cell exhaustion to the tumor and subsequently trapping them, driving them towards residence^35,36^. Given the exclusive presence of 4-1BB^+^ T cells in the tumor (Figure 2B) and that protection against metastases would require circulation of T cells, we hypothesized that treatment with neoadjuvant anti-4-1BB induces the egress of antigen-specific T cells from the primary tumor to pre-metastatic sites prior to surgical resection, thereby mediating systemic protection. Surprisingly, we noted the highest baseline expression of resident marker CD103 in T_prog_ (Figure S6A, B). We reasoned that CD103 plays an important role in T_prog_, with alternative molecules such as CXCR6 serving the role of T cell retention in T_int_ and T_term_ subsets^71,79^. Focusing on T_prog_, we noted that 6-day window (6D), but not 3-day window (3D), anti-4-1BB drove a significant reduction in CD103 in the E0771.lmb model (Figure 4A). Importantly, this reduction in CD103 was mirrored in the 4T1 model (Figure S6C). Taken together, these data suggest given sufficient time, neoadjuvant anti-4-1BB drives T_ex_ out of tumors through downregulation of resident markers.

**Figure 4.**
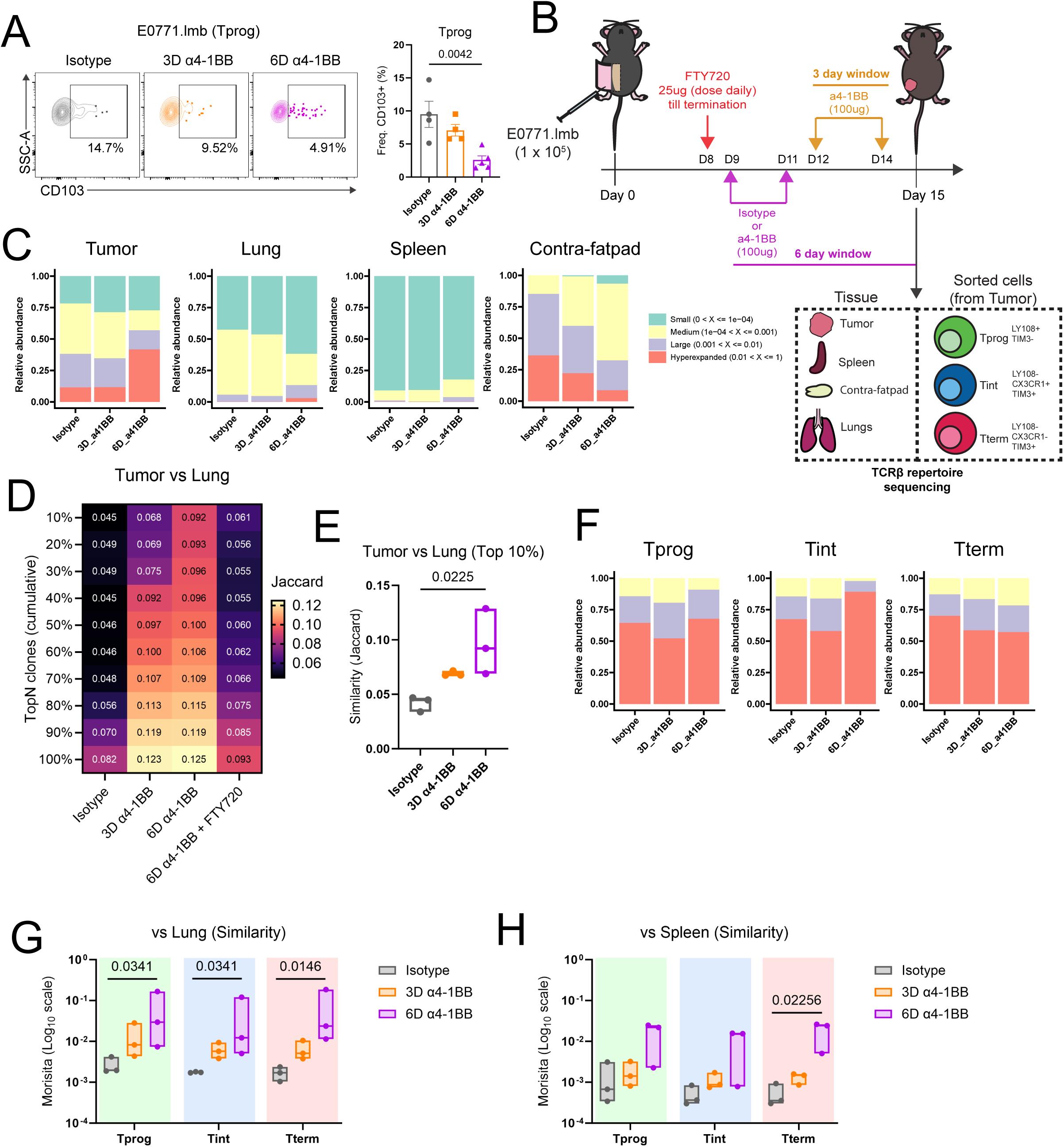
Anti-4-1BB drives reduction of resident marker in intratumor T cells, and concomitant increased clonotype overlap with distant tissues. **(A)** Representative flow plot showing CD103^+^ levels of T_prog_ treated with isotype, 3D α4-1BB (3-day window) or 6D α4-1BB (6-day window) in E0771.lmb (left), quantification of CD103^+^ frequencies (right). (Isotype, n = 4; 3D α4-1BB, n = 4; 6D α4-1BB, n = 5). **(B)** Experimental design for bulk TCR sequencing of E0771.lmb model. **(C)** Stacked Barplot illustrating relative abundance of T cells in tumor, lung, spleen and contra-lateral fatpad, grouped by proportional abundance. **(D)** Heatmap showing T cell similarity between tumor and lung by Jaccard similarity index. TCRs ranked and cumulatively binned by abundance. **(E)** T cell similarity of top 10% clonotypes between tumor and lung. **(F)** Stacked Barplot illustrating relative abundance of sorted T_prog_, T_int_ and T_term_, grouped by proportional abundance**. (G-H)** Morisita overlap index comparing sorted cells and lung (G) or spleen (H) harvested from mice under treatments with isotype, 3D α4-1BB or 6D α4-1BB. (in B-H, Isotype, n = 3; 3D α4-1BB, n = 3; 6D α4-1BB, n = 3, 6D α4-1BB + FTY720, n = 3). Data are presented as mean ± SEM. *P* values were determined using one-way ANOVA followed by post-hoc Dunnett’s multiple comparisons test (A), Kruskal-Wallis test followed by post-hoc Dunn’s multiple comparisons test (E, G, H).

To assess how different treatment windows of anti-4-1BB influenced the overall structure of TCR repertoire across tissue compartments, we performed bulk TCRβ repertoire sequencing on E0771.lmb tumors, lungs, contralateral fatpad and spleen, without T cell purification using our previously established protocol^71,100–103^ (Figure 4B). We noted that 6D but not 3D window anti-4-1BB, drove clonal expansion of T cells within the primary tumor (Figure 4C). Importantly, this effect was not seen in the other tissues. To determine the degree of unique TCR clonotype overlap between the primary tumor obtained from independent mice, we calculated the Jaccard index as previously described^71,104^. By ranking and binning TCRs within each tissue by clonal abundance, we noted a significant increase in similarity of TCR in the top 10% clones detected, suggesting preferential expansion of similar clonotypes across mice (Figure S7A). In agreement, we noted a moderate, albeit not significant, decrease in diversity of TCR in 6D anti-4-1BB treated tumor as assessed by Shannon entropy (Figure S7B). Together, these results confirm that the primary response to anti-4-1BB occurs within the primary tumor.

Next, to determine if the T cells expanded by anti-4-1BB in the primary tumor could egress into circulation and colonize distant tissues, we compared the degree of clonotype sharing between primary tumor and distant tissues. Clonotype sharing between tumor and lungs increased in anti-4-1BB relative to isotype treated groups, with a correlation between the extent of overlap and window of treatment (Figure 4D). In line with our egress hypothesis, this form of clonotype sharing was brought back to isotype levels upon FTY720 treatment. Importantly, we noted a significant increase in similarity of TCR clonotypes in the top 10% clones detected in tumor and lung following 6D anti-4-1BB treatment relative to isotype (Figure 4E). In addition, the same trend was held when comparing tumor clonotype sharing with spleen, albeit not reaching statistical significance (Figure S7C). To directly account for both clonotype amount and abundance among the shared clones, we used the Morisita index^71,105^. The Morisita index revealed a trend of anti-4-1BB in increasing TCR sharing between tumor to lung, but not to spleen and contralateral fatpad (Figure S7D). Together, these results suggest that given sufficient time, anti-4-1BB drives preferential egress of T cells from the tumor to lung, reinforcing the importance of the length of window between anti-4-1BB treatment to surgical resection.

Given that we noticed a decline in residence marker in T_prog_ after anti-4-1BB treatment (Figure 4A), we sought to validate if there was preferential egress of select T_ex_ subsets following anti-4-1BB treatment. To do so, we sorted T_prog_, T_int_, and T_term_ from E0771.lmb tumors in parallel with whole tissues (Figure 4B). 6D anti-4-1BB drove the highest level of clonal expansion in T_int_ relative to T_prog_ and T_term_ while neither 3D anti-4-1BB nor isotype treatment resulted in noticeable clonal expansion (Figure 4F). The finding is in line with preferential expression of 4-1BB on T_int_ relative to other T_ex_ subsets (Figure 2M-O), suggesting higher levels of 4-1BB correlate with increased responsiveness to anti-4-1BB. However, using the Morisita index to assess global T cell similarity between the sorted cells and lungs, we found that treatment with 6D anti-4-1BB resulted in significant increase in similarity across all 3 T_ex_ subsets (Figure 4G). This result is not surprising given the lineage relationship between the 3 T_ex_ subsets and their shared TCRs (Figure S7E, F). Importantly, we did not observe a significant increase in sharing for majority of subsets to neither the spleen nor contralateral fatpad (Figure 4H, S7G), suggesting preferential tissue conditions in the lung to allow seeding of T_ex_ originating from the tumor.

### Expansion of T_int_ correlates with shared clonotypes at distant tissue under optimal anti-4-1BB treatment window

Given the lineage relationship between the three T_ex_ subsets (Figure S7E, F), it is possible anti-4-1BB preferentially impacted a specific subset which then propagated across subsets. To investigate this possibility, we first ranked and binned the top 100 TCRs of each individual sorted T cell subset and visualized their frequencies in the lung. We found that the top 20 clones of each subset were consistently expanded in the 6D anti-4-1BB relative to both 3D anti-4-1BB and isotype-treated groups (Figure 5A). Focusing on the top 20 TCRs, we sought to determine how many were shared amongst the subsets within the respective treatment groups. To this end, we computed and visualized the distinct clonotypes shared among the three subsets, between any two subsets, or within a single subset (Figure 5B-D). Despite the dominant number of shared TCRs among the T_ex_ subsets, we found that only frequencies of T_int_, and not T_prog_ or T_term_, treated under the 6D anti-4-1BB regimen correlated significantly with the frequencies of T cells with the shared TCRs in either the lung or spleen (Figure 5E-G, S8). This indicated that T_int_ is the primary responder to anti-4-1BB though we cannot exclude the contribution from the other T_ex_ subsets. Taken together, our data indicates that the administration of anti-4-1BB promotes preferential expansion of T_int_ which colonizes, suggesting usage in the neoadjuvant setting may serve to fortify metastases-prone sites such as the lungs.

**Figure 5.**
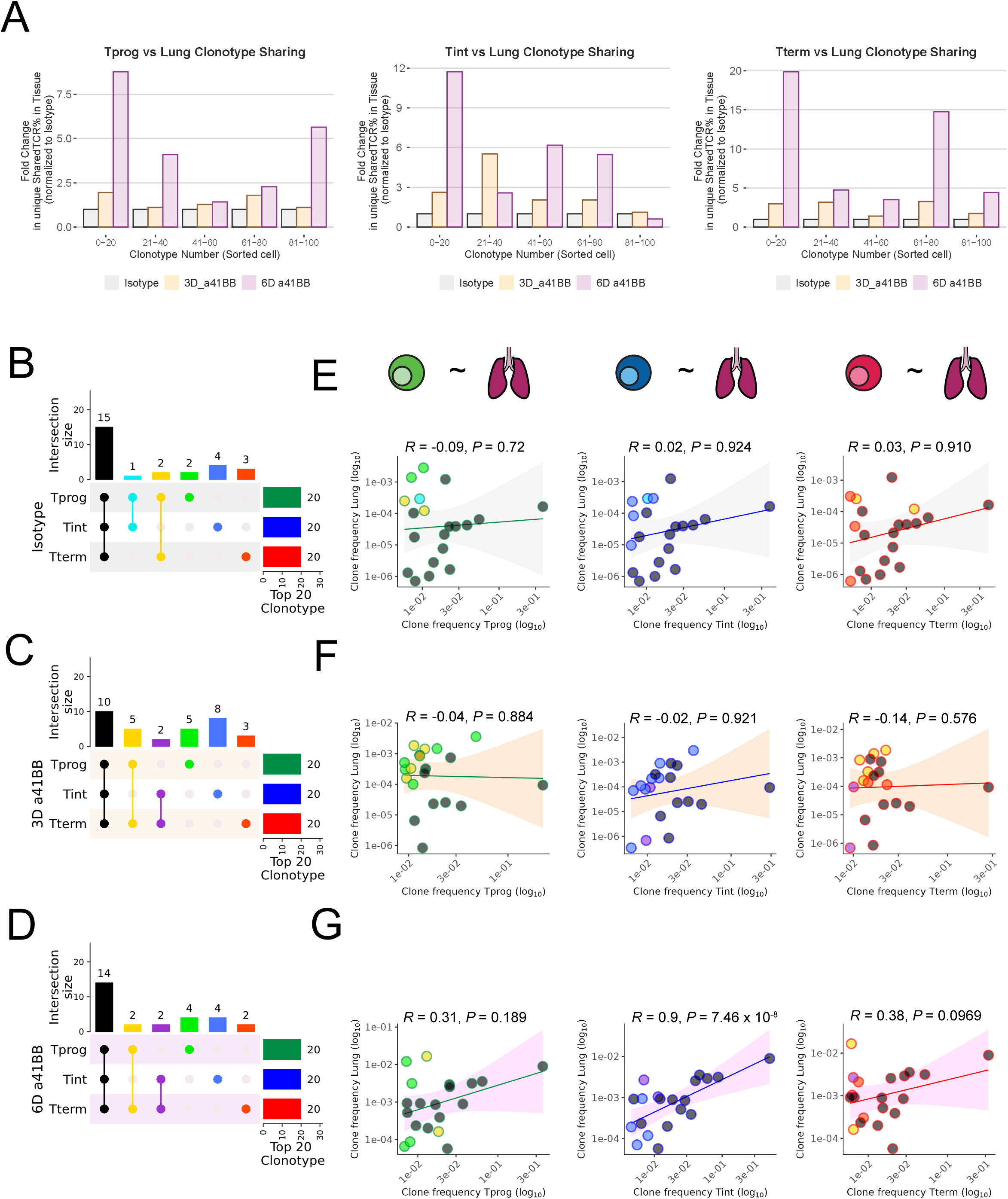
Significant correlation between expanded clonotype in T_int_ and distant tissue. **(A)** Barplot depicting fold change in TCR frequency in the lung following treatment with 3D anti-4-1BB or 6D anti-4-1BB. Frequencies are normalized isotype frequencies in respective bins. TCRs are first identified from sorted cells, ranked and binned by abundance within the sorted cell groups. **(B-D)** UpSet plots showing number of distinct TCR clonotypes belonging to each combination of T_ex_ subset, and not found in other combinations, among the top 20 clonotypes in either isotype (B), 3D anti-4-1BB (C) or 6D anti-4-1BB treated groups (D). For clarity, bar and numbers at the top enumerates the number of clonotypes shared in the specific combination of T_ex_ subsets highlighted. (Black =All; Cyan = Tprog and Tint only; Yellow = Tprog and Tterm only; light Green = Tprog only; light Blue = Tint only; light red = Tterm only. Bar and numbers on the right shows the 20 unique clonotypes from Tprog (dark Green), Tint (dark Blue) and Tterm (dark Red) used in the comparisons. **(E-G)** Scatter plots comparing the frequencies of clonotypes in Tprog (left), Tint (middle), Tterm (right) to respective frequencies in lung under Isotype (E), 3D anti-4-1BB (F) or 6D anti-4-1BB (G) treatment. Correlation coefficient calculated using *cor.test* function. Pearson’s *R* and *P* value was determined using a two-sided t distribution with n − 2 degrees of freedom. Shaded area represents 95% confidence interval of linear model as determined by *geom_smooth* function (grey = Isotype; orange = 3D anti-4-1BB; pink = 6D anti-4-1BB).

### Anti-4-1BB signature is predictive of BRCA patient prognosis

While anti-4-1BB has not entered clinical trials in the neoadjuvant setting, we sought to evaluate the prognostic significance of T_ex_ treated with anti-4-1BB. To do so, we conducted single-cell RNA sequencing (scRNA-seq) on sorted CD45^+^ immune cells from E0771.lmb tumors treated with isotype or anti-4-1BB under a 3-day window as a part of a broader study (Figure S9A). Tumor-infiltrating CD8^+^ T cells clustered distinctly from splenocytes (Figure S9B). We extracted tumor-infiltrating CD8^+^ T cells, re-clustered, and projected onto UMAP to obtain 10 clusters (Figure 6A). Beyond visualizing the expression of the most variable genes (Figure S9C), we annotated the clusters by overlaying transcriptional signatures of multiple T_ex_ subsets, obtained from a published study^106^ utilizing the chronic LCMV infection model, onto our UMAP (Figure 6B, S9D).

**Figure 6.**
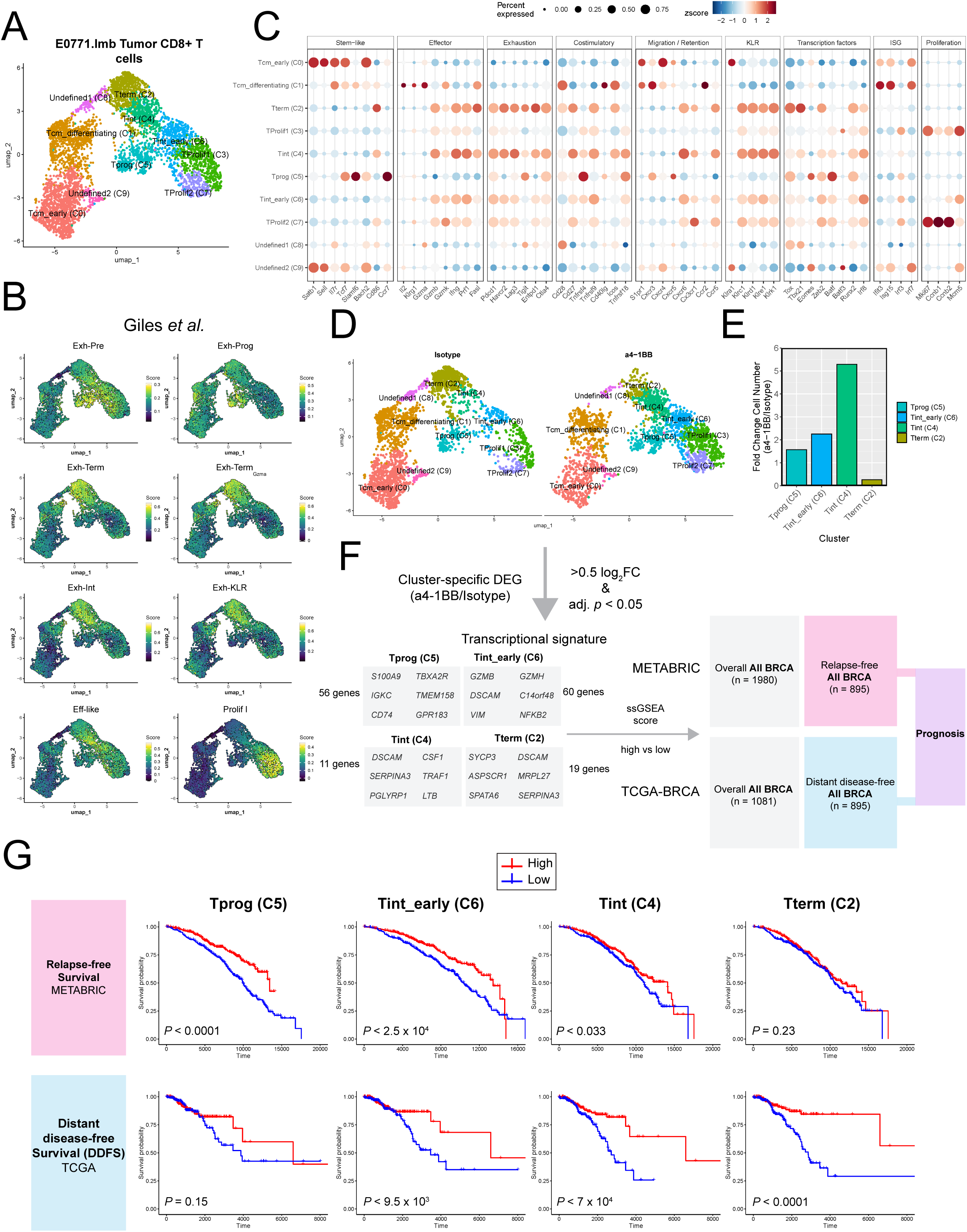
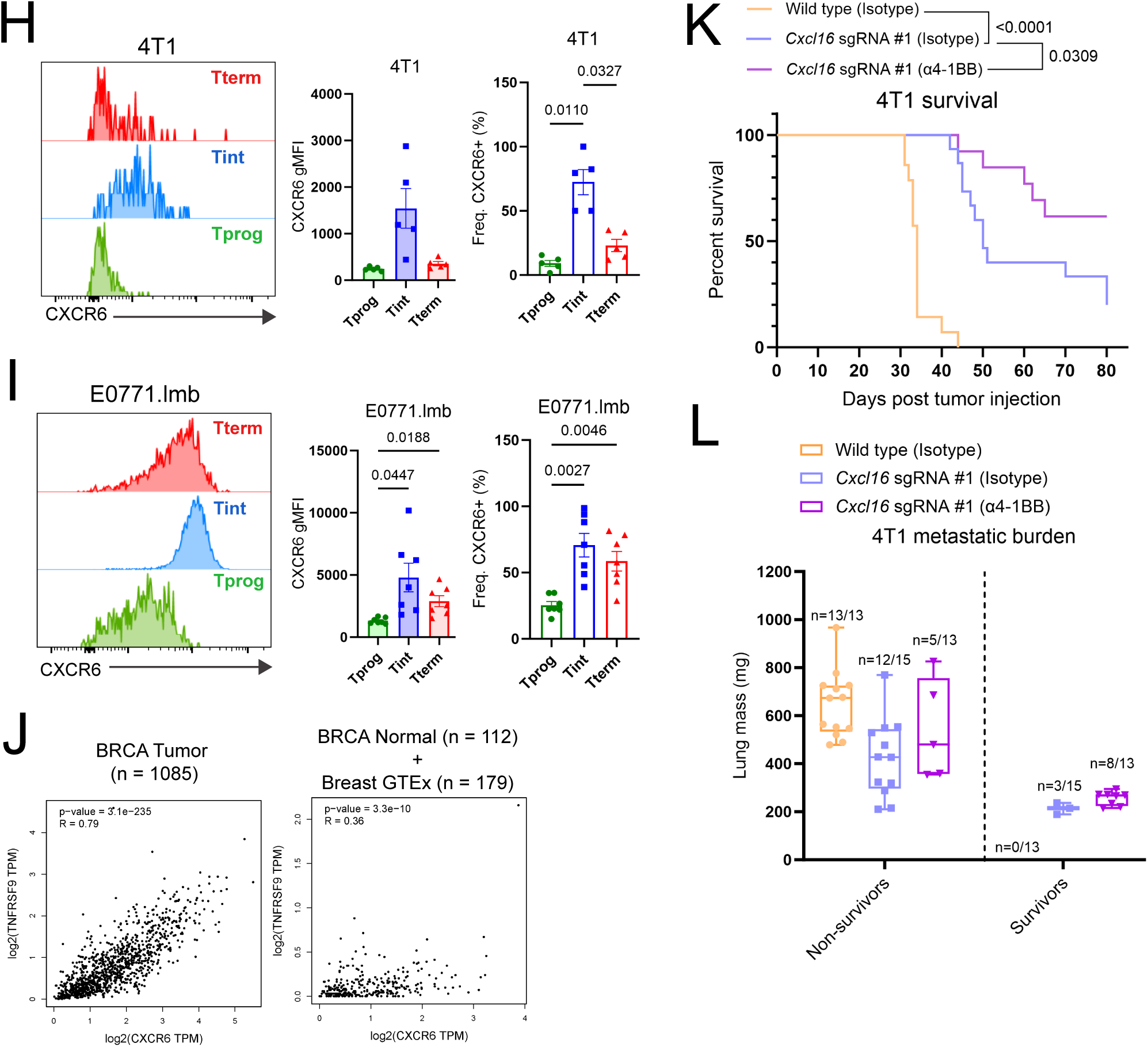
Anti-4-1BB predicts relapse-free survival in human BRCA patients and synergizes with loss of CXCL16. **(A)** UMAP of total CD8^+^ T cells from E0771.lmb tumors. **(B)** Enrichment of T_ex_ subset specific gene signatures from gp33-specific T cells from chronic LCMV infection model (clone 13) isolated on day 15 and 30 (Giles *et al.*). **(C)** Dotplot showing expression of stem-like, effector, exhaustion, costimulatory, chemokine receptor, killer cell lectin-like receptor (KLR), transcription factors, interferon-stimulated genes (ISG) and proliferation-related genes across clusters. **(D)** UMAP CD8^+^ T cells split by treatment, isotype (left), anti-4-1BB (right). **(E)** Ratio of specific T_ex_ subsets following anti-4-1BB treatment. **(F)** Schematic diagram depicting workflow for generating T_ex_ subset specific anti-4-1BB augmented transcriptional signatures from E0771.lmb scRNA-seq dataset and applying signatures to BRCA patient specimen data from TCGA and METABRIC. **(G)** Total BRCA Relapse-free survival from METABRIC study (top) and Distant-Disease free relapse survival from TCGA (bottom) after separating high (3^rd^ quantile) and low (1^st^ quantile) gene signature expression groups. **(H-I)** Representative flow plots showing higher CXCR6 expression in T_int_ relative to T_prog_ and T_term_ (left), gMFI of CXCR6 (middle), frequency of CXCR6^+^ (right), from models 4T1 (H), E0771.lmb (I). **(J)** Correlation between CXCR6 and TNFRSF9 expression in human TGCA BRCA tumor (left) and TCGA BRCA normal and breast GTEx samples (right). Spearman’s rank-correlation coefficient r and associated *p*-value are shown. Analysis done on GEPIA2 webtool. **(K)** Kaplan-Meier analysis of 4T1 wild-type or *Cxcl16* sgRNA #1 treated with isotype or anti-4-1BB on days 13 and 15 post-tumor injection. Primary tumors were surgically resected on day 17 and mice underwent survival analysis. **(L)** Lung metastatic burden (mass) at endpoints. (in K-L, wild type (Isotype), n = 13-14; *Cxcl16* sgRNA #1 (isotype), n = 15; *Cxcl16* sgRNA #1 (α4-1BB), n = 13). *P* values were determined using log-rank (Mantel-Cox) test (G, K), one-way ANOVA followed by post-hoc Tukey’s multiple comparisons test (H, I).

We observed cluster 0 – early central memory T cells (T_cm_early_; *Sell, Tcf7*, *Satb1*), cluster 1 – differentiating central memory T cells (T_cm_differentiating_; *Il7r*, *Cd28*, *Gzma*, *Ifit3*, *Isg15*), clusters 3 and 7 – proliferating T cells (T_Prolif_; *Mki67^high^*, *Ccnb2, Mcm5*), cluster 6 – early intermediate exhausted T cells (T_int_early_; *Irf8*, *Spp1*), cluster 5 – progenitor exhausted T cells (T_prog_; *Slamf6*, *Ccr7*, *Cd83*), cluster 4 – intermediated exhausted T cells (T_int_; *Tnfrsf9^high^*, *Cxcr6^high^*, *Gzmb*, *Ifng*), cluster 2 – terminally exhausted T cells (T_term;_ *Havcr2*, *Entpd1, Tox^high^*), clusters 8 and 9 – undefined cells (Figure 6C, S9D). Focusing only on T_ex_ subsets that express high levels of *Pdcd1* (clusters 2 to 7), we performed pseudotime analyses by placing T_prog_ (cluster 5) as the starting point. Indeed, pseudotime analyses confirmed a branch point T_int_early_ (cluster 6) which subsequently developed into T_int_ and T_term_ on one branch or T_Prolif1/2_ on another branch (Figure S9E).

By splitting the UMAP by treatment (Figure 6D), we noticed that anti-4-1BB increases T_prog_, T_int early_ and T_int_ while decreasing in T_term_ clusters (Figure 6E). This suggests that there is potential reinvigoration of CD8^+^ TILs when treated with anti-4-1BB. To evaluate the potential prognostic significance of anti-4-1BB treatment, a gene signature specific for each cluster was generated based on the differentially expressed genes (DEGs) detected comparing anti-4-1BB to isotype for respective clusters and overlaid onto human BRCA datasets from METABRIC and TCGA (Figure 6F). Focusing on relapse-free survival and distant disease-free relapse survival from METABRIC and TCGA BRCA respectively, we noted that signature obtained from T_int_early_ and T_int_ consistently correlated with positive prognosis across both datasets (Figure 6G). Signatures obtained from T_prog_ and T_term_ were only associated with positive prognosis in recurrence-free survival either one of the datasets (Figure 6G). Interestingly, we noticed this prognostic effect afforded by anti-4-1BB-enhanced T_int_early_ and T_int_ signatures was largely diminished in overall survival in the METABRIC dataset, supporting our hypothesis that 4-1BB signaling correlates with reduced recurrence-survival (Figure S9D). Taken together, this suggests that 4-1BB signaling within the primary tumor can counteract subsequent recurrence.

### Anti-4-1BB synergizes with loss of tumor-derived CXCL16 to counteract metastases in CD8^+^ T cell-dependent manner

While we found that anti-4-1BB can promote the expansion and egress of T cell from the primary tumor into pre-metastatic sites, it is constrained by the narrow window of opportunity. Given the potential complications arising from extending the window of neoadjuvant treatment schedule in the clinic, we sought to determine potential targets with the TME that could be targeted to synergize with anti-4-1BB treatment. In tandem with the theme of T cell retention, our group previously demonstrated that CXCL16, the sole ligand for CXCR6, traps CXCR6^+^ T_EFF/EM_ within the primary tumor of breast cancer and that these T cells have anti-metastatic function that can be harnessed when released into peripheral sites with intra-tumoral administration of anti-CXCL16 prior to surgical resection^71^. Importantly, CXCL16 is enriched in basal subtype of breast cancer in humans, relative to normal tissues (Figure S10A). As such, we sought to determine if ablation of CXCL16 solely from tumor cells would be sufficient to mediate protection against metastasis. We generated *Cxcl16*KO lines on the 4T1 background using CRISPR/Cas9 (Figure S10B-E). We isolated 2 clones which showed a moderately different proliferation rate from wild type (WT) 4T1 in vitro (Figure S10F). When injected in vivo, both clones presented slower primary tumor growth as opposed to WT (Figure S10G-H). More importantly, both clones demonstrated lower metastatic burden and prolonged survival as opposed to WT (Figure S10I-J), recapitulating our previous findings using the antibody blockade^71^.

Previous publications have indicated that T_term_ but not T_prog_ express CXCR6 across multiple tumor types^79^. We further interrogated and demonstrated that T_int_ and T_term_ but not T_prog_ express CXCR6 (Figure 6H, I). Importantly, analysis of human breast cancer data shows a strong correlation between CXCR6 and 4-1BB (Figure 6J). Using the 4-day treatment window schedule – which was insufficient in achieve long-term term survival in the context of neoadjuvant anti-4-1BB monotherapy (Figure S2E) – we sought to determine if anti-4-1BB could synergize with the loss of tumor-derived CXCL16 to mediate long-term survival (Figure S11A). Anti-4-1BB treatment on *Cxcl16*KO 4T1 tumors resulted in long-term survivors under the 4-day window treatment schedule (Figure 6K, J). Furthermore, by depleting CD8^+^ T cells through the administration of anti-CD8β, we demonstrate that the survival benefit observed in *Cxcl16*KO tumors, with or without anti-4-1BB treatment, is in part dependent on CD8^+^ T cells (Figure S11B-F). Together, these findings suggest that blocking tumor-derived CXCL16 in CXCL16^high^ TNBC could be a potential complement to the effects of neoadjuvant anti-4-1BB on released T_ex_, thereby further fortifying distant tissues against metastases.

## DISCUSSION

Neoadjuvant immunotherapy is a promising avenue to achieve durable control over metastatic disease^22^. However, widespread adoption of checkpoint blockade in treating early-stage TNBC has been limited due to heterogeneity of the disease^65^. More recently, studies have pointed to the importance of 4-1BB/4-1BBL signaling axis in licensing checkpoint blockade^107,108^. In this study, we sought to reorient the emphasis to the use of anti-4-1BB as a monotherapy to uncover the mechanism underpinning its success within the neoadjuvant setting in preclinical models. Exhausted TIL subsets comprise of a heterogeneous population of CD8^+^ T cells^26^. Here, we identified that T_int_, and more broadly low-affinity T cells, most highly expresses 4-1BB across multiple TNBC models (Figure 2K-M), which we pose to be the major responders to anti-4-1BB. Indeed, given an optimal scheduling window, TCRβ sequencing revealed T_int_ to be major responders to anti-4-1BB – evidently through clonal expansion (Figure 4F). Importantly, circulating T cells that expanded within the primary tumor have been demonstrated to reflect the functions of their tumor-infiltrating counterparts^109^.

Many cancer immunology studies utilize primary tumor growth as a readout for treatment efficacy^110,111^, insofar as a surrogate for metastatic cancers with tumor dissemination, which is the main cause of mortality in the clinic^112–114^. Because cancer is a systemic disease, it requires a systemic approach to understand and assess patient outcomes after treatment^45,115^. To demonstrate the clinical relevance of our findings, we drew upon our scRNA-seq dataset, deriving cluster-specific gene signatures following anti-4-1BB treatment (Figure 6G). We demonstrated the prognostic significance of anti-4-1BB treated T_int_ signatures when applied to both TCGA and METABRIC BRCA datasets – remarkably on distant-disease. Notably, this would be indicative of endogenous 4-1BB/4-1BBL signaling given that BRCA patients are, to-date, not treated with anti-4-1BB. Given that APC-rich niches within the TME have been shown to provide the necessary costimulatory signals needed sustain CD8^+^ TILs for productive anti-tumor immunity and response to ICB^78,108,116,117^, our finding that 4-1BB enriched gene signature in specific TIL subsets being prognostic in a subset of BRCA patients would not be hyperbolic.

Protection against metastatic recurrence has been attributed to the functions of memory T cells^52^. The affinity of TCR repertoire for an antigen is important in driving a diverse recall response to mediate protection^51^. To substantiate this, our group has recently demonstrated the importance of low-affinity T cells in counteracting tumor relapse in the context of combination ICB therapies^102,103^. Conversely, high-affinity TCR has been reported to increase T cell retention through establishment of tissue residency program within non-lymphoid tissue and tumor while leading to rapid dysfunction^90,99,118^. To counteract retention, our group and others have previously demonstrated how driving the egress of T cells out of the tumor prior to surgical resection leads to better systemic protection^41,42,71,119^. Here, we extended this notion beyond conventional ICB approaches. We demonstrated that, with optimal scheduling, anti-4-1BB monotherapy can downregulate resident marker CD103 and drive the egress of multiple T_ex_ subsets into pre-metastatic tissues. Furthermore, we show that weak TCR affinity enriched for the T_int_ subset, and more broadly, 4-1BB expression (Figure 3I, J). Given that low-affinity T cells are canonically associated with memory generation^51^, and that anti-4-1BB enhances memory T cell formation^120–122^, it is likely that neoadjuvant anti-4-1BB drives the egress of low-affinity memory precursor T cells into circulation which subsequently fortifies pre-metastatic tissues. Further studies would be needed to validate this hypothesis. Finally, our findings that the combination of anti-4-1BB and reduction of CXCL16 from the TME resulted in a synergistic effect and sustained survival suggests this combination is a potential candidate for future implementation in the clinic under the neoadjuvant setting.

In this era of cancer immunology, the prognosis for TNBC remains dismal. Major immunotherapeutic efforts are focused on the development of personalized therapies involving adoptive transfer of engineered T cells^123^. Notably, the endodomain of 4-1BB was adopted since the inception of second-generation chimeric antigen receptor T cell, demonstrating superior capacity in counteracting exhaustion as compared to that of CD28^124^. Nevertheless, these engineered T cell modalities are limited by manufacturing costs, deep prior knowledge and have shown limited success in solid tumors^125,126^. Neoadjuvant immunotherapy offers a broad, and cost-effective approach, inducing robust systemic immune response by exploiting the presence of the primary tumor and its endogenous tumor-reactive T cell pool. Preclinical studies have identified immune suppressive niches within TNBC that hypoxic in nature, thereby resisting T cell attack^127^. Furthermore, the hypoxic microenvironment was previously shown to contribute to upregulation of 4-1BB expression in TILs^128^. Our findings highlight anti-4-1BB as a clinically relevant treatment option in the neoadjuvant setting for TNBC, whose administration presents three novel advantages: 1) low-affinity intratumor T cells express high levels of 4-1BB, 2) anti-4-1BB can drive the clonal expansion of specific intratumor T cell subsets that are relatively malleable and 3) become inclined to exit the tumor, escaping the immune-subversive tumor microenvironment prior to irreversible exhaustion, subsequently seeding distant tissue sites to mediate protection. In this way, we hope to eliminate the source, while curbing the spread. While the concept of promoting T cells egress from the primary tumor may run counterintuitive to anti-tumor immune response, an increasing number of publications recognize the need and importance to consider dissemination of antigen-reactive T cells into systemic circulation to achieve long-term remission in the face of metastatic disease.

## Materials and Methods

### Mice

All mouse experiments were performed with approval from the Duke University Animal Care and Use Committee. C57BL/6 (Strain: 000664), TcraKO (Strain: 002116), OT-2 (Strain: 004194) and BALB/cJ (Strain: 000651) were purchased from The Jackson Laboratory. Rag2-KO BALB/c mice (C.129S6(B6)-Rag2tm1Fwa N12) were purchased from Taconic. OT-1; Thy1.1 transgenic mice were generated by crossing OT-1 (Strain: 003831) and B6 Thy1.1 (Strain: 000406). Only female mice aged 7-12 weeks were used for all the experiments. Mice were housed under pathogen-free conditions at 22°C, with a humidity between 30% and 70%, a light/dark cycle of 12 h and ad libitum access to food and water.

### Cell culture

4T1 (CRL-2539) and HEK293T cells (CRL-3216) were obtained from Duke Cell Culture facility with original sourcing from ATCC. E0771 (CRL-3461), E0771.lmb (CRL-3405) and Py8119 (CRL-3278) were obtained directly from ATCC. 4T1 was cultured in high glucose RPMI 1640, supplemented with 10% fetal bovine serum, 1% penicillin/streptomycin, 1mM sodium pyruvate, and 10mM HEPES. E0771 and E0771.lmb were cultured in Dulbecco’s modified Eagle’s medium (DMEM) supplemented with 10% fetal bovine serum, 1% penicillin/streptomycin and 20mM HEPES. HEK293T was cultured in Dulbecco’s modified Eagle’s medium (DMEM) supplemented with 10% fetal bovine serum and 1% penicillin/streptomycin. Py8119 was cultured in Ham’s F-12K (Kaighn’s) media supplemented with 5% fetal bovine serum and 1% penicillin/streptomycin. Mouse T cells were cultured in RPMI 1640 media supplemented with 10% fetal bovine serum, 1% penicillin/streptomycin, 2mM L-glutamine, 1mM sodium pyruvate, 0.1mM nonessential amino acids, and 50μM 2-mercaptoethanol. All cells were cultured in a humidified 37°C incubator with 5% CO_2_.

### Stable cell line generation

HEK293T cells with approximately 50-70% confluency was transfected with transfer plasmids together with lentiviral packaging vectors psPAX2 (Addgene: 12260) and VSV.G (Addgene: 14888) using Lipofectamine 2000 (Thermo Fisher Scientific, 11668019). Lentiviral particles were collected 48h post-transfection and filtered through 0.45µm PES filter. For transduction, viral supernatant was added to cultured cells with 8 μg/mL Polybrene (Sigma-Aldrich, TR-1003). 24h post-transduction, viral supernatant was replaced with fresh media.

To generate tumor OVA APL overexpression cell lines, OVA (N4) or its APL (Q4, T4, V4) was cloned onto pWPT vector (Addgene: 12255). eGFP was replaced with either mScarlet-I or mTurquoise2 (obtained from Addgene: 98839) as indicated in experimental schematics. A GSG linker preceding the T2A self-cleaving peptide sequence was introduced inbetween the OVA APL and reporter to enhance cleavage efficiency. In summary, the following tumor lines were generated:

Py8119-OVA (N4)-T2A-eGFP
Py8119-OVA (Q4)-T2A-Scarlet
Py8119-OVA (T4)-T2A-Turquoise
E0771-OVA (N4)-T2A-eGFP
E0771.lmb-OVA (N4)-T2A-eGFP
E0771.lmb-OVA (Q4)-T2A-eGFP
E0771.lmb-OVA (T4)-T2A-eGFP

After transduction, reporter positive cells were enriched by FACs for 2 to 3 rounds to achieve purity of at least 90%.

### CRISPR-Cas9 editing

To generate 4T1 *Cxcl16KO* cell lines, single guide RNA sequences targeting exon 2 of mouse *Cxcl16* was designed using Benchling (https://www.benchling.com/), selecting for gRNA with ON-target score > 50 and OFF-target score > 50. The sequences of sgRNA used in this study are:

*Cxcl16* sgRNA #1: 5’-ATGTGATCCAAAGTACCCTG-3’
*Cxcl16* sgRNA #2: 5’-TCTGGCACCCAGATACCGCA-3’

Complementary guide oligonucleotides were annealed and cloned into the BsmBI site of the lentiCRISPR v2 (Addgene: 52961) plasmid according to the Zhang laboratory’s protocol^129^. 4T1 cells were seeded at 50,000 cells per well in 12-well plates. In less than 12 hours, 4T1 was transiently transfected using 2 µg of gRNA in lentiCRISPR v2 to Lipofectamine 2000 in a 1:3 ratio (µg:µl). After 12 hours of overnight incubation, cells were split into new 12-well plates at a ratio of 1:10 in complete media. The following day, media was changed to complete media with the addition of 2 µg/ml of puromycin. The selection process was carried out for 3 days. Clones were then obtained using limiting dilution on a 96-well plate. These wells were expanded and validated for knockout by western blot and flow cytometry.

### Western blot

4T1 cells were seeded at a density of an estimated 100,000 cells per well in 6-well plates. In less than 24 hours post-seeding, media of cells was changed to complete media supplemented with either vehicle treatment or 20ng/ml IFNγ (Peprotech) plus 1µM GI254023X (Sigma Aldrich). Cells were harvested 24 hours post-treated and lysed using 30ul RIPA Buffer (Sigma) plus 1:100 of Halt Protease and Phosphatase inhibitor cocktail (Thermo Fisher Scientific). Protein concentrations were measured using Pierce BCA protein assay kit (Thermo Fisher Scientific). Equal amounts of protein were then loaded into Mini-PROTEAN TGX Stain-Free Precast Gels (Biorad #4568083). Proteins were transferred to PVDF membrane (Thermo Fisher Scientific #88520) and blocked in 5% normal donkey serum (v/v) (Jackson ImmunoResearch Lab Inc., #017-000-121) in 1X TBST. Membranes were probed with primary Polyclonal Goat IgG anti-mouse Cxcl16 (R&D Systems AF503) at 1:2,000 dilution from 0.2 µg/ml stock, and primary Rabbit anti-mouse β-actin (Cell Signaling Technology #4967) at 1:1,000 dilution from stock concentration overnight at 4 ℃. Next, membranes were probed for secondary IRDye® 800CW Donkey anti-Goat IgG (LI-COR #926-32214) at 1:15,000 dilution and secondary Alex Fluor 680-conjugated Goat Anti-Rabbit IgG (H+L) (Jackson ImmunoResearch Lab Inc. #111-625-144) at 1:5,000 dilution for Cxcl16 and β-actin respectively. Membranes were imaged using the LI-COR Odyssey Infrared Imaging System and inverted using ImageStudio Lite version 5.2.

### In vitro proliferation assay (MTT)

We performed in vitro proliferation assay using RealTime-Glo™ MT Cell Viability Assay following manufacturer’s instructions. Briefly, cells are seeded at 5000 cells per well and luminescence readouts taken every 2 or 4 hours for 48 hours. Relative light units (RLU) are normalized to readout taken at the first hour after seeding of cells.

### Mouse tumor models

For 4T1, E0771.lmb, E0771, Py8119 – including all derivatives of respective lines obtained through knockout or overexpression methodologies – tumor cells were harvested and resuspend in PBS. Indicated number of cells used for inoculation is stated in individual experimental schematic of respective figures. Concentration of cells was adjusted to desired dose per 50µl of PBS. Using a 27-gauge needle, tumor cells were orthotopically implanted into the fourth mammary fatpad of mice under anesthesia with isoflurane. Tumor volume was monitored every 3 days post tumor injection (PTI) or indicated otherwise, until tumor resection or if it reached a maximum of 2000mm^3^ in volume. Tumor volume (mm^3^) = 0.5 × length (mm) × [width (mm)]^2^.

### Immunotherapy and in vivo lymphocyte depletion

For anti-4-1BB therapy, tumor-bearing mice were intraperitoneally injected with either 100µg of anti-mouse 4-1BB (3H3) antibodies (Bio X Cell, BE0239) or IgG2a isotype control (Bio X Cell, BE0089) at indicated days. For anti-PD-1 therapy, tumor-bearing mice were injected with either anti-mouse PD-1 (J43) antibodies (Bio X Cell, BE0033-2) or IgG isotype control (Bio X Cell, BE0091) at indicated dose on indicated days. For lymphocyte depletion experiments, mice were intraperitoneally injected with anti-mouse CD8β (53-5.8) antibodies (Bio X Cell, BE0223) or IgG1 isotype control (Bio X Cell, BE0088) every 4 days after tumor injection until surgical resection, and every 7 days post-surgical resection.

### Adoptive T cell transfer

For adoptive transfer of OT-1 and OT-2 cells into tumor-bearing mice, lymph nodes and spleen were harvested from respective transgenic mice and mechanically dissociated. Red blood cells were lysed with ammonium-chloride-potassium (ACK) lysis buffer (Quality Biological). Naïve OT-1 and OT-2 were then negatively selected, and isolated using Dynabeads Untouched mouse CD8 (Invitrogen 11417D) and CD4 (Invitrogen 11415D) kits respectively. T cells were then mixed, resuspended in PBS and intravenously injected via the tail vein route.

### Surgical resection and mouse survival studies

Prior to surgical resection of tumors, mice were anaesthetized with isoflurane and injected subcutaneously with 5mg/kg of meloxicam for pain management. Curved dissecting scissors and blunt forceps were used to surgically remove tumors. Sterile clips (Stoelting) were used to close wounds following surgery and mice were placed on warm heat pad until recovery was ensured with additional supportive care provided. For survival phase of study, mice were monitored until humane endpoint was reached, indicated by weight loss of 15 to 20%, lethargy, breathing difficulties or moribund status. At individual endpoint of mice, lungs were harvested, weighed and subsequently stored in Bouin solution for visualization and enumeration of metastatic nodules. Either lung mass or lung tumor nodules are used as metastatic burden readout as indicated in plots. For lung metastatic nodule counts, we counted nodules three times per lung and averaged the counts. Mice that experienced local tumor recurrence were excluded from the survival study and lung metastatic burden readouts.

### Flow cytometry

Flow cytometry for CXCL16 on 4T1 tumor cells in vitro was conducted as previously described^71^. To investigate tumor-infiltrating lymphocytes, mouse breast tumor tissues were harvested and draining lymph node (inguinal) was carefully separated from tumor. Tumors were mechanically dissociated followed by further enzymatic digestion by incubating samples in 1mg/ml of collegenase IV (Gibco) in HBSS supplemented with 100µg/ml DNase1 (Roche) and 50mM MgCl_2_ for 30-45 min at 37°C. Red blood cells were removed by incubation with ACK lysis buffer and subsequent washing with PBS. Lymph nodes were only mechanically dissociated. Samples were filtered through 70µm nylon mesh. Single cell suspensions were first stained with LIVE/DEAD™ Fixable Aqua Dead Cell Stain Kit (Invitrogen) at 1:1000 in PBS. For cell surface marker staining, samples resuspended in FACs buffer (PBS supplemented with 2% FBS and 2mM EDTA) and treated with Fc receptor blocking antibody (Bio X Cell, 2.4G2) for 10 min at 4°C followed by surface antibodies (1:100-200 dilution) for another 20 min at 4°C. For intracellular staining, eBioscience™ Foxp3/Transcription Factor Staining Buffer Set was used to fix and permeabilize cells according to manufacturer’s instructions. To assess the absolute number of lymphocytes in tumor tissues, absolute counting beads (Invitrogen) were added into each sample. Samples were acquired with a BD FACSCanto II or LSRFortessa X-20 machine (BD Biosciences), and data were analyzed using FlowJo software v10.

### Antibodies for flow cytometry

Invitrogen

**Anti-mouse CD8α (KT15)**: FITC
Biolegend

**Anti-mouse CD45.2 (104):** APC/Cy7
**Anti-mouse TCRβ (H57-597):** APC/Cy7
**Anti-mouse CD8α (53-6.7):** BV605
**Anti-mouse CD62L (MEL-14):** PE/Dazzle™ 594
**Anti-mouse CD103 (2E7):** BV421, BV785
**Anti-mouse CXCR4 (L276F12):** PE/Dazzle™ 594
**Anti-mouse CXCR6 (SA051D1)**: BV711
**Anti-mouse CX3CR1 (SA011F11):** FITC, BV421, BV785
**Anti-mouse PD-1 (RMP1-30)**: PE/Cy7
**Anti-mouse 4-1BB (17B5):** PE, APC
**Anti-mouse TIM3 (RMT3-23):** APC, BV785
**Anti-mouse LY108 (330-AJ)**: APC
**Anti-mouse LAT2 (NAP70):** Alexa Fluor 647 BD
Bioscience

**Anti-mouse CD44 (IM7):** BUV395
**Anti-mouse CD8α (53-6.7):** BUV737
**Anti-mouse CD69 (H1.2F3):** BUV737
**Anti-mouse PD-1 (RMP1-30):** BUV738
**Anti-mouse TCF1 (S33-966)**: PE
**Anti-rat/mouse CD90.1 (OX-7):** BV786
CST

**Anti-human/mouse TCF1 (C63D9):** Alexa Fluor 488

### Tetramer binding analysis

For mouse tetramer binding analysis, cells were pretreated with the Fc receptor blocking antibody (Flow cytometry method). Cells were then stained with H2-K^b^ SIINFEKL tetramer BV421 (NIH Tetramer Core Facility, Atlanta, GA, USA) at a final concentration of 6µg/ml for 30 min at 4°C. This was followed by surface staining and flow cytometry analysis.

### TCRβ repertoire sequencing

1 × 10^5^ E0771.lmb tumors orthotopically implanted into the mammary fat pad of female B6 mice. For 6-day window, mice were administered i.p. with 100 µg of anti-mouse 4-1BB (BioXCell, #BE0239) or isotype control (BioXCell, #BE0089) on days 9 and 11. For 3-day window, mice were administered i.p. with 100 µg of anti-mouse 4-1BB (BioXCell, #BE0239) on days 9 and 11. 25 µg of FT720 or PBS was administered on day 8 daily until endpoint of study. Primary tumor, lung, contralateral fatpad and spleen were isolated from E0771.lmb tumor-bearing mice 15 days post-tumor injection from all treatment groups and lysed in TRI Reagent^®^ solution (Sigma Aldrich) using lysis beads (Zymo Research). In parallel, T_prog_, T_int_ and T_term_ were sorted from primary tumors directly into TRI Reagent^®^ solution, using the Beckman-Coulter Astrios sorter. Samples were spiked with 2B4.11 cells as a normalization control as previously reported^130^. RNA was extracted using the Direct-zol RNA kit (Zymo Research) according to the manufacturer’s instructions. 1µg or lower amount of RNA was converted to cDNA using the qScriptFlex cDNA synthesis kit (QuantaBio) with constant region-specific primer. Multiplex PCR was performed to amplify the CDR3 region of rearranged TCRβ loci. A set of primers, each specific to a specific TCR Vβ segments, and a reverse primer to the constant region of TCRβ were used to generate a library of amplicons that cover the entire CDR3 region. PCR products were loaded onto 2.5% agarose gels and bands between 300-350 bp were excised and purified using the GeneJET Gel Extraction kit (ThermoFisher). These purified products were sequenced using the Illumina NovaSeq X platform (Novogene).

### TCRβ repertoire analyses

For analyses of bulk TCRβ-seq, raw sequencing data was preprocessed and aligned to TCR gene segments using the MiXCR software^131^. Numbers of distinct TCR clonotypes and accumulative clone frequencies (both amino acid and nucleotide sequences) were calculated after normalization using in-house R scripts. The Immunarch package was used for clonotype summary, repertoire diversity, and similarity analysis. Venn diagrams were visualized using VennDiagram package (v1.7.3). UpSet plots were made with UpSetR package (v.1.4.0) using the *make_comb_mat* function in ‘distinct’ mode to obtain distinct clonotypes within each combination of groups.

### scRNA sequencing

2.5 ×10^5^ E0771.lmb tumors were orthotopically implanted into the mammary fat pad of 2-month-old female B6 mice. On days 14 and 16, mice were administered i.p. with 100 µg of anti-mouse 4-1BB (BioXCell, #BE0239) or isotype control (BioXCell, #BE0089). On day 17, Live CD45^+^ cells were sorted from the tumors and spleens of 4 mice per group, and 20,000 cells were loaded onto the Chromium Next GEM Chip (10x Genomics) to target for 10,000 recovered cells per sample. The library preparation was conducted using the Single Cell 3’ v3.1 chemistry following the manufacturer’s instructions (10x Genomics). Briefly, the first-stranded cDNA was generated in the Gel Beads-in-emulsion (GEMs), magnetically purified after the collapse of GEMs and amplified via PCR. Enzymatic fragmentation and addition of sample indexes were then conducted for the construction of sequencing libraries. Paired-ended sequencing was performed on the NovaSeq 6000 system (Illumina).

### scRNA-seq analyses

For analyses of 4T1 scRNA-seq, “TumorTem_counts” file was sourced from https://data.mendeley.com/datasets/compare/3f4rsk96kf/4. The gene count matrix was subjected to quality control, pre-processing and clustering using R package Seurat^132^ (version 5.0.3). Cells with total number of genes expressed (nFeature_RNA) between 200 and 3000 were kept. Next, we excluded cells whose mitochondrial gene counts accounted for more than 10% of total counts. Expression data were then normalized, scaled, and searched for variable features using the *SCTransform* v2 function^133^. We used a resolution of 0.5 for the *FindCluster* function to group cells. We subsequently kept only CD8^+^ T cells and repeated the clustering process with the default parameters for the aforementioned functions for further analysis used in this manuscript.

For analyses of E0771.lmb scRNA-seq, an aggregate count matrix was generated using Cell Ranger (version 7.1.0, 10x Genomics). Briefly, reads were aligned to the mm10 genome using the cellranger count pipeline. The cellranger aggr was then used to normalize counts across samples. Next, we processed the aggregated matrix using the R package Seurat (version 4.2.1). Cells with a total number of genes (nFeature_RNA) between 200 and 7500 and a gene count (nCount_RNA) lower than 60,000 were kept. We further excluded cells whose mitochondrial gene counts accounted for more than 10% of total counts. Highly variable genes were detected using *FindVariableFeatures* function with the nFeatures parameter set to 2000. PCA was first implemented, and the top 15 PCs were used for UMAP transformation and cluster analysis. After annotation, only CD8^+^ T cells from tumor samples were kept for subsequent analyses in this manuscript. Subsequent analysis for E0771.lmb was performed on R package Seurat (version 5.0.3). We performed *ScaleData* using all genes. And finally, we used the *FindCluster* function at resolution 0.8. For both 4T1 and E0771.lmb scRNA-seq, uniform manifold approximation and projection was used for dimension reduction and visualization. Pseudotime trajectory analyses was performed using Monocle3^134^ using Seurat wrapper functions *cluster_cells*, *learn_graph*, *order_cells*. For obtaining DEGs, we used Seurat *FindMarkers* function, keeping only cells that fulfilled the criteria of log2 fold change > 0.5, adjusted *p* value < 0.05, and expressed on more than 10% of cells in cluster. For projection of published gene signatures on E0771.lmb UMAP, DEG from Supplementary table 1 from Giles *et al.* was obtained, passed into Seurat *AddModuleScore* function, and projected onto the UMAP. UMAP of signature score was generated using SCpubr^135^. *CalcStats* in SeuratExtend^136^ was used to calculate z-score for variable gene expression between clusters. Violin plot for signature score and heatmap showing variable gene expression were generated using SeuratExtend. Otherwise, native Seurat functions were used to generate plots.

### Gene signature and Human BRCA survival analyses

The RNA-Seq normalized (TPM) gene expression profiles for the TCGA-BRCA dataset, along with corresponding phenotype information and sample survival data, were obtained from the UCSC Xena database (https://xenabrowser.net/datapages/). Samples lacking survival data or derived from non-tumor tissues were excluded from further analysis. METABRIC breast cancer data were downloaded from the cBioPortal platform^137^. To examine the role of 4-1BB signaling across the different T_ex_ subsets as a potential prognostic factor, we derived cluster specific gene signatures from the E0771.lmb scRNA-seq data. To obtain DEGs, we used Seurat *FindMarkers* function, keeping only cells that fulfilled the criteria of log_2_ fold change > 0.5, adjusted *p* value < 0.05, and expressed on more than 10% of cells in individual clusters. To obtain total counts in TCGA BRCA dataset, only samples with the “01A” suffix (tumor tissues) were retained (n = 1081). For obtaining total counts METABRIC dataset, all samples with gene expression profiles were included (n = 1980). For application of gene signature to obtain recurrence free survival prognosis, we used the “Relapse-free” (n = 1176) and ‘Distant Disease-free” (n = 895) metrics for the METABRIC and TCGA BRCA datasets respectively. DEGs (anti-4-1BB over isotype) of respective clusters were mapped to orthologous genes using the Ensembl database using the bioMart package^138^. Mapped genes were used for calculation of enrichment score using the Single-sample Gene Set Enrichment Analysis (ssGSEA)^139^ algorithm implemented in package, Gene Set Variation Analysis for Microarray and RNA-Seq Data (GSVA) version 2.0.4. Samples were classified based on ssGSEA score using upper and lower quantile methods and Kaplan-Meier survival analysis was then performed using survminer package to evaluate the correlation between feature gene sets and patient survival outcomes.

### Statistics and Reproducibility

No statistical methods were used to predetermine sample sizes, but our sample sizes are similar to those reported in previous publications. Data distribution was assumed to be normal, but this was not formally tested. For each mouse experiment, mice were randomly assigned to each group, based on their age and weight, to eliminate the age and weight differences. Prior to treatment, mice were allocated based on tumor volume so that each group had an equitable starting point in terms of tumor volume. Data collection and analysis were not performed blind to the conditions of the experiments. Flow cytometry data of tumors were excluded if contaminated with dLN. Survival and lung metastatic burden data was excluded if there was local primary tumor recurrence. Statistical analysis was performed using GraphPad Prism 10 software (GraphPad Software). Two-way analysis of variance (ANOVA) was used to compare tumor growth curves, taking the interaction term as the statistical readout. Survival benefits were evaluated by Kaplan–Meier methods, and log-rank (Mantel-Cox) test was used to calculate statistical significance. Two-tailed Student’s t-test or Wilcox test was performed to compare statistical difference between two groups, as indicated. For tumor mass or flow cytometry data, one-way ANOVA was used to compare outcomes across multiple experimental groups. For multiple comparisons, Dunnett, Tukey or Šídák’s corrections were used to adjust *P*-values as indicated in figure legend. For index comparisons, Kruskal-Wallis test followed by post-hoc Dunn’s multiple comparisons test. Differences with a minimum of *P* < 0.05 were considered as statistically significant. For correlation analysis of top 20 clones of sorted cells frequencies to respective frequencies in lung or spleen, the correlation coefficient calculated using *cor.test* function. Pearson’s R and P value was determined using a two-sided t distribution with n − 2 degrees of freedom. For correlation analysis of human *CXCR6* and *TNFRSF9* and comparing expression of *CXCL16*, TCGA/GTEx data available within GEPIA2^140^ (http://gepia2.cancer-pku.cn/) was used to directly to generate the plots. For correlation analysis of human *CXCR6* and *TNFRSF9* expression, correlation coefficient (*R*) and *p*-value were obtained using Spearman correlation analysis. For comparison of *CXCL16* expressing between Tumor and Normal tissues, the “Expression DIY” tab was used. Data are first log_2_(TPM+1) transformed for differential analysis with log_2_FC is defined as median(Tumor) - median(Normal). Statistical comparison is made using one-way ANOVA.

## Acknowledgements

We thank the Lynn Martinek, Kayla Parr and Javid Mohammed of the Duke University Flow Cytometry Shared Resource and Andrew Chan of the Flow Cytometry Platform at Singapore Immunology Network (SIgN), A*STAR for assisting with cell sorting. We thank the staff of Duke University’s Division of Laboratory Animal Resources for supporting and accommodating our study schedules. We thank Shanshan Wu Howland, Alicia Tay and Ivy Foo of the Immunogenomics Platform at Singapore Immunology Network (SIgN), A*STAR for assisting with the scRNA sequencing. We thank the NIH Tetramer Core Facility for providing tetramer reagents. We thank all members of Wang and Li lab, past and present, for the valuable discussions and contributions to this project. This work was supported by the NUS Development Grant to B.J.W.L. This work was supported by R01-CA233205 and R01-CA249726 from the NIH to X.-F.W. and Q.-J.L. Q.-J.L. is supported by core research grants provided to IMCB and SIgN by the Biomedical Research Council (BMRC), A*STAR and National Research Foundation (NRF) Singapore under the NRF Investigatorship (NRFI09-0016).

## Author contributions

B.J.W.L., X.-F.W. and Q.-J.L. conceptualized the study. B.J.W.L performed the majority of the experiments. M.L. and F.P.-L.T. performed the scRNA sequencing. B.J.W.L., M.L. and L.W. analyzed the scRNA-seq data. B.J.W.L. and S.L.Y.K. performed the TCRβ sequencing. L.W. analyzed all TCRβ-seq data. T.Y. assisted with surgeries. T.Y., C.Y. and K.X. contributed to the in vivo experimental setup. C.C. and H.W. performed the projection of gene signatures to human survival datasets. A.M. assisted with flow cytometry. E.W., Q.J., Z.M., L.T., R.N.C., D.Q. and C.C.P assisted with experiments and analysis. B.J.W.L. wrote the manuscript. X.-F.W. and Q.-J.L. revised the manuscript. X.-F.W. and Q.-J.L. secured the funding, reagents and oversaw the study.

## Supplementary Figures

**Supplementary Figure 1.**
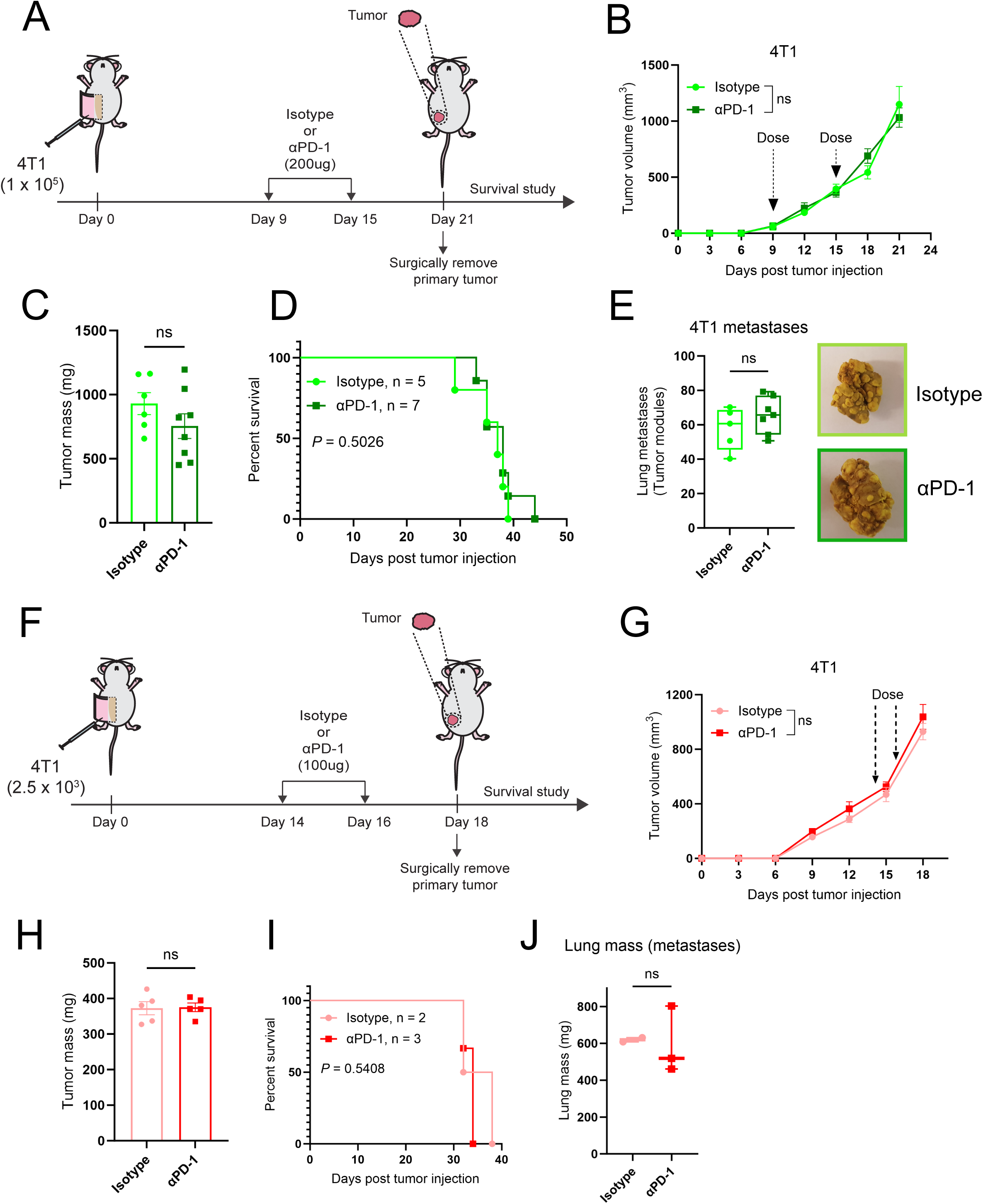
Neoadjuvant anti-PD-1 does not protect against spontaneous metastasis. – related to Figure 1. **(A-E**) 11**-**day window anti-PD-1 experimental timeline for orthotopic 4T1 metastasis model. **(A)** Experimental schematic. **(B)** Primary tumor growth. **(C)** Primary tumor mass. **(D)** Kaplan-Meier analysis. **(E)** Lung metastatic nodule counts at endpoints, enumeration (left) and representative picture (right) (in B-C, Isotype, n = 7; α4-1BB, n = 8; in D-E, Isotype, n = 5; α4-1BB, n = 7). **(F-J**) 4**-**day window anti-PD-1 experimental timeline for orthotopic 4T1 metastasis model. **(F)** Experimental schematic. **(G)** Primary tumor growth. **(H)** Primary tumor mass. **(I)** Kaplan-Meier analysis. **(J)** Lung metastatic burden (mass) at endpoints (in G-H, Isotype, n = 5; α4-1BB, n = 5; in I-J, Isotype, n = 2; α4-1BB, n = 3). Data are presented as mean ± SEM. *P* values were determined using two-way repeated measures ANOVA (B, G), two-tailed unpaired Student’s *t*-test (C, E, H, J), log-rank (Mantel-Cox) test (D, I); ns, not significant.

**Supplementary Figure 2.**
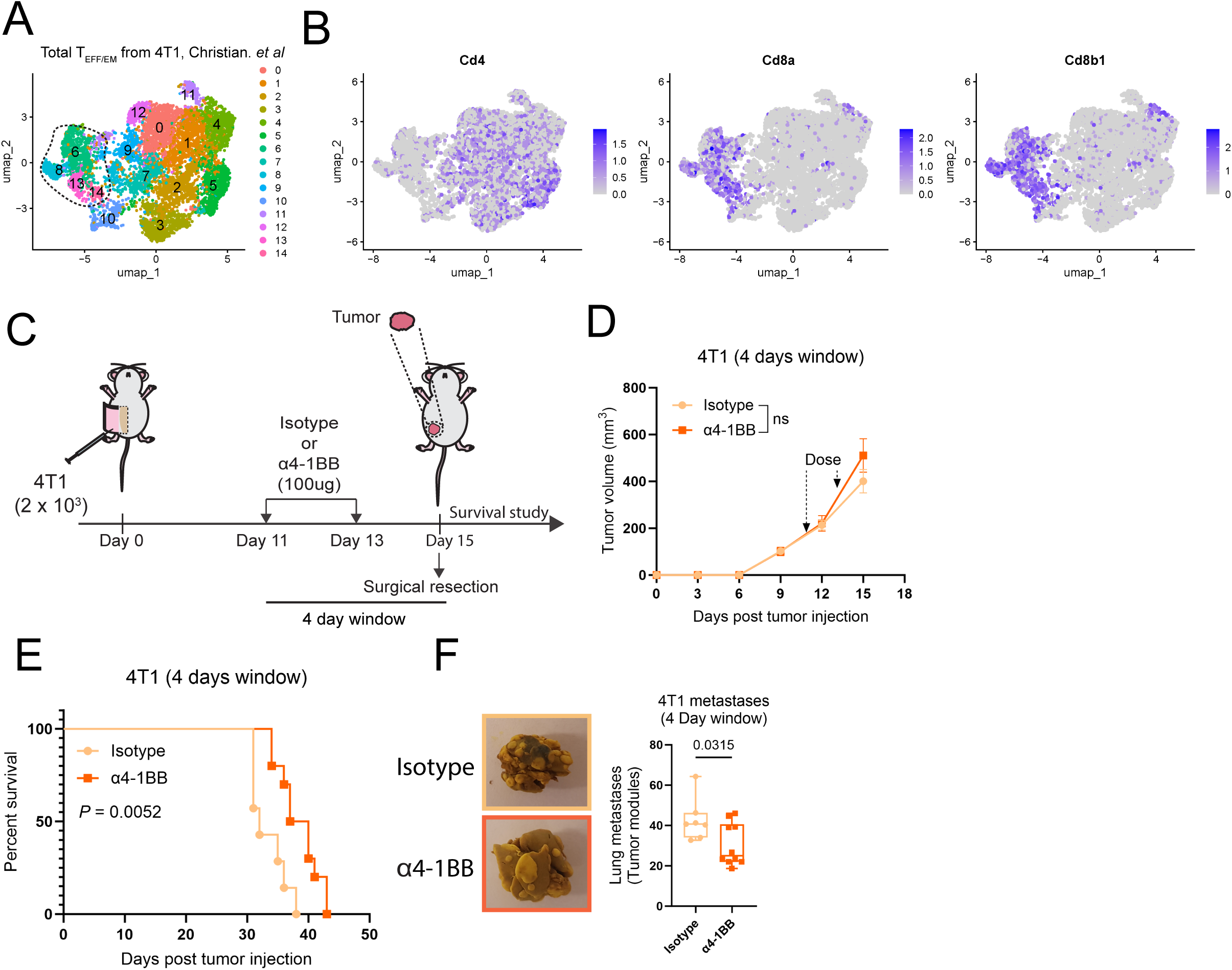
Neoadjuvant anti-4-1BB prolongs survival in spontaneous metastasis murine model. – related to Figure 1. **(A)** UMAP of total T_EFF/EM_ clusters from *Christian et al.* (B) Relative *Cd4*, *Cd8a* and *Cd8b1* levels. Clusters 6, 8, 13 and 14 (traced by dashed lines) are taken for subsequent CD8^+^ T_EFF/EM_ only analysis. **(C-F**) 4**-**day window experimental timeline for orthotopic 4T1 metastasis model. **(C)** Experimental schematic. **(D)** Primary tumor growth. **(E)** Kaplan-Meier analysis. **(F)** Lung metastatic nodule counts at endpoints, representative picture (left) and enumeration (right) (in D, Isotype, n = 11; α4-1BB, n = 12; in E, Isotype, n = 7; α4-1BB, n = 10. Data is pooled from two independent experiments). *P* values were determined using two-way repeated measures ANOVA (D), log-rank (Mantel-Cox) test (E), two-tailed unpaired Student’s *t*-test (F); ns, not significant.

**Supplementary Figure 3.**
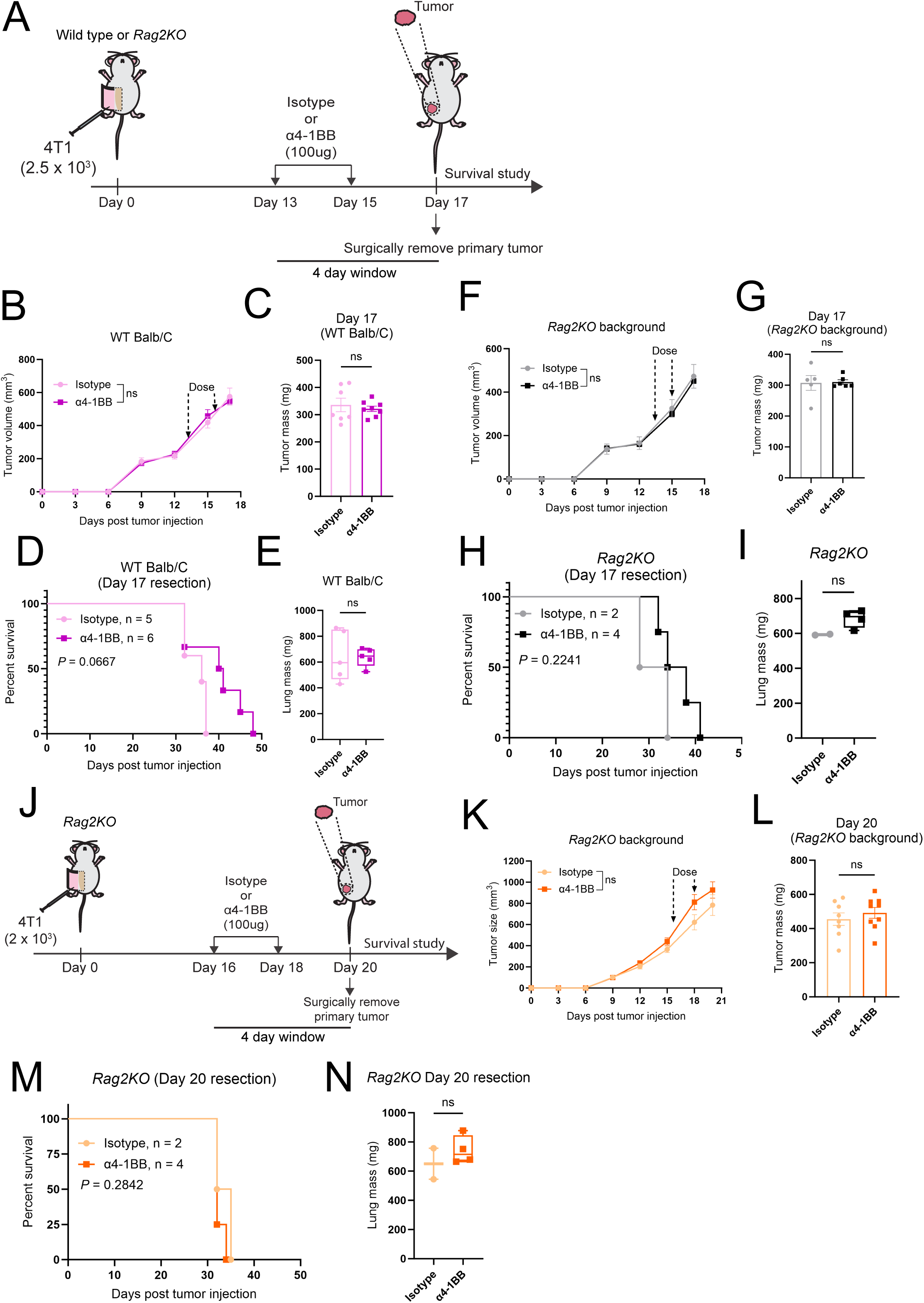
Anti-4-1BB mediated survival benefit is dependent on T cells – related to Figure 1. **(A)** 4**-**day window experimental timeline for orthotopic 4T1 metastasis model on Day 17 resection in wild type mice (B-E), or *Rag2KO* mice (F-I). **(B)** Primary tumor growth. **(C)** Primary tumor mass. **(D)** Kaplan-Meier analysis. **(E)** Lung metastatic burden (mass) at endpoints (in B-C, Isotype, n = 7; α4-1BB, n = 8; in D-E, Isotype, n = 5; α4-1BB, n = 6)**. (F)** Primary tumor growth. **(G)** Primary tumor mass. **(H)** Kaplan-Meier analysis. **(I)** Lung metastatic burden (mass) at endpoints (in F-G, Isotype, n = 5; α4-1BB, n = 6; in H-I, Isotype, n = 2; α4-1BB, n = 4)**. (J-N**) 4**-**day window experimental timeline for orthotopic 4T1 metastasis model on Day 20 resection in *Rag2KO* mice. **(J)** Experimental schematic. **(K)** Primary tumor growth. **(L)** Primary tumor mass. **(M)** Kaplan-Meier analysis. **(N)** Lung metastatic burden (mass) at endpoints (in K-L, Isotype, n = 8; α4-1BB, n = 9; in M-N, Isotype, n = 2; α4-1BB, n = 4.). Data are presented as mean ± SEM. *P* values were determined using two-way repeated measures ANOVA (B, F, K), two-tailed unpaired Student’s *t*-test (C, E, G, I, L, N), log-rank (Mantel-Cox) test (D, H, M); ns, not significant.

**Supplementary Figure 4.**
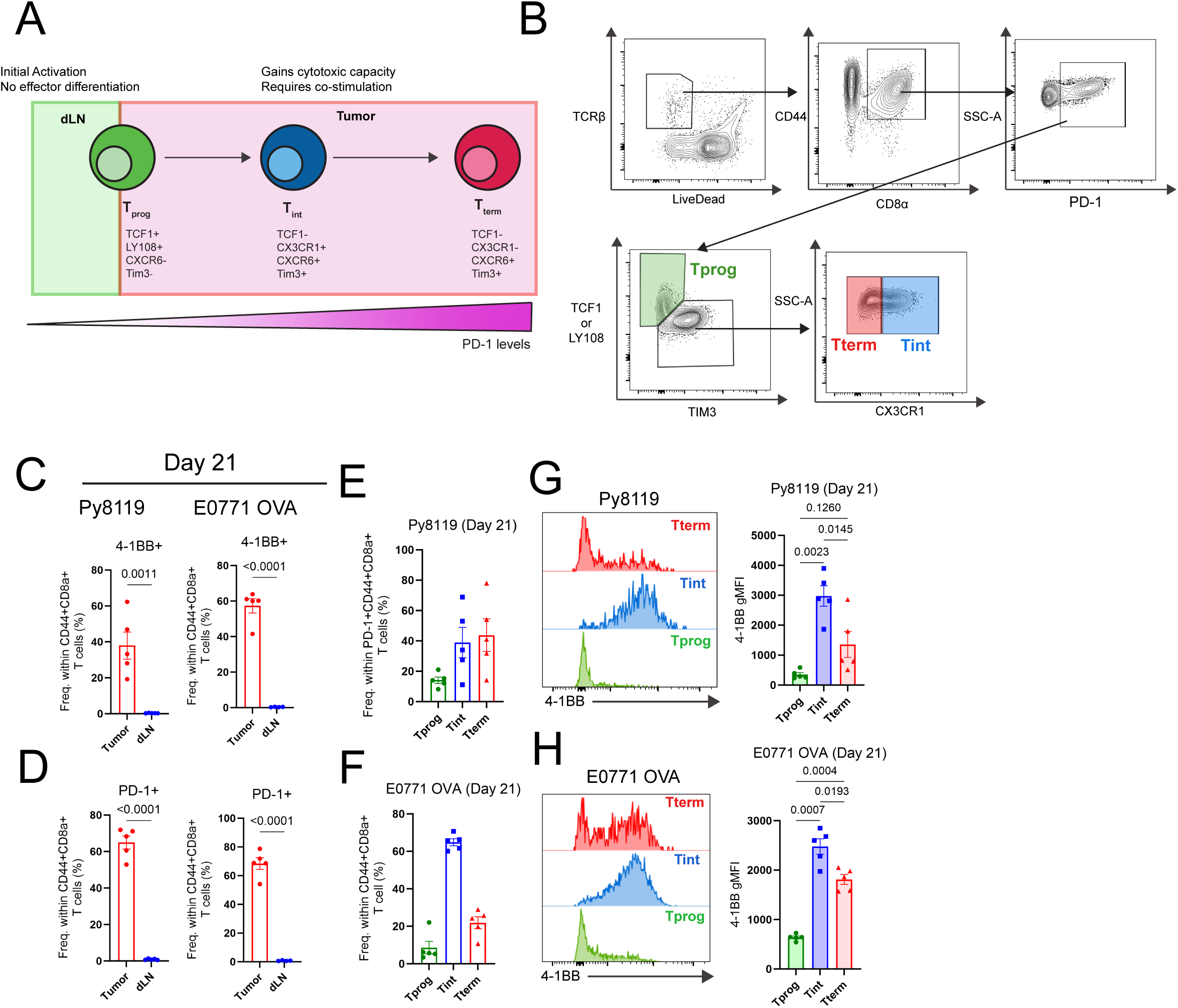
Enrichment of 4-1BB only in TILs and not dLN – related to Figure 2. **(A)** Model of Tex subsets by tissue location, depicting transition of early T_prog_ entrants from dLN into tumor and subsequent seeding of T_int_ and T_term_ in that order. **(B)** Representative gating strategy for T_prog_, T_int_ and T_term_ in tumor samples (shown here is E0771.lmb tumor). **(C)** Frequency of 4-1BB in CD44^+^CD8^+^ T cells of tumor and dLN of Py8119 (left) and E0771 OVA (right) tumor models at Day 21. **(D)** Frequency of PD-1 in CD44^+^CD8^+^ T cells of tumor and dLN of Py8119 (left) and E0771 OVA (right) tumor models at Day 21. **(E, F)** Frequency of T_prog_, T_int_ and T_term_ in TILs of Py8119 (E) and E0771 OVA (F) tumors at Day 21. **(G)** Representative flow plots showing 4-1BB expression (left) and gMFI (right) of 4-1BB in T_prog_, T_int_ and T_term_ in Py8119 at Day 21. **(H)** Representative flow plots showing 4-1BB expression (left) and gMFI (right) of 4-1BB in T_prog_, T_int_ and T_term_ in E0771 OVA at Day 21. (in C-H, Py8119, n = 5; E0771 OVA, n = 5). Data are presented as mean ± SEM. *P* values were determined using two-tailed unpaired Student’s *t*-test (C, D), one-way ANOVA followed by post-hoc Tukey’s multiple comparisons test (G, H).

**Supplementary Figure 5.**
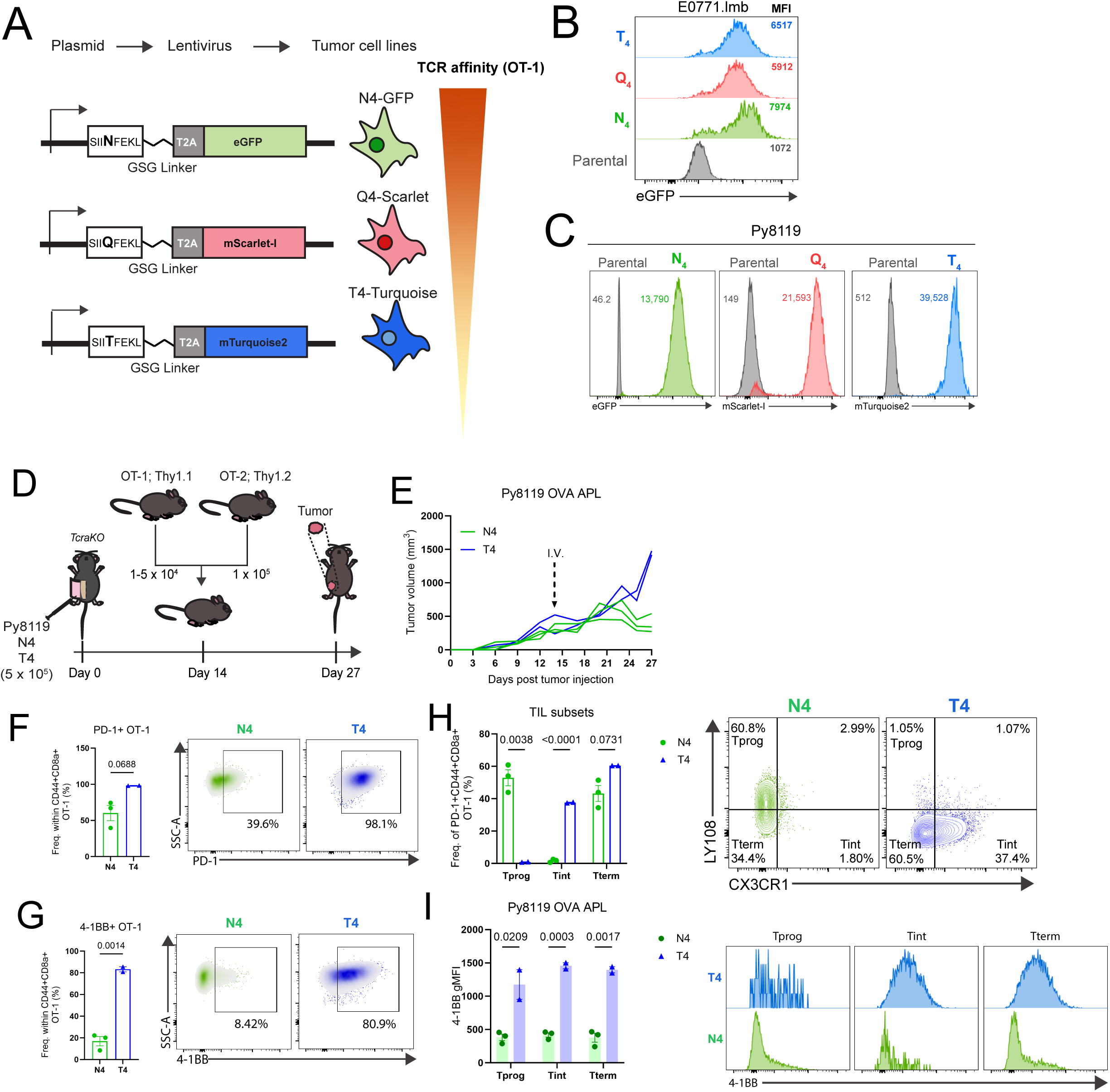
Generation and validation of in vivo tumor OVA APL system. – related to Figure 3. **(A)** Generation of OVA APL system. **(B)** Expression of eGFP in E0771.lmb as surrogate for OVA APL expression. **(C)** Expression of respective reporter proteins in Py8119 as surrogate for OVA APL expression. **(D-I)** T_int_ is enriched in low-affinity T cells in Py8119. **(D)** Experimental schematic. **(E)** Tumor volume of Py8119 expressing OVA APL following transfer of OT-1 and OT-2. **(F)** Frequency (left) and representative FACs plots (right) of PD-1^+^ OT-1. **(G)** Frequency (left) and representative FACs plots (right) of 4-1BB^+^ OT-1. **(H)** Frequency of T_prog_, T_int_ and T_term_ in PD-1^+^ compartment of OT-1 (left), representative flow plots (right). **(I)** gMFI of 4-1BB in T_prog_, T_int_ and T_term_ (left). Representative flow plots showing 4-1BB expression (right) (in F-J, n = 2-3). Data are presented as mean ± SEM. *P* values were determined using two-way repeated measures ANOVA (E), two-tailed unpaired Student’s *t*-test (F-I).

**Supplementary Figure 6.**
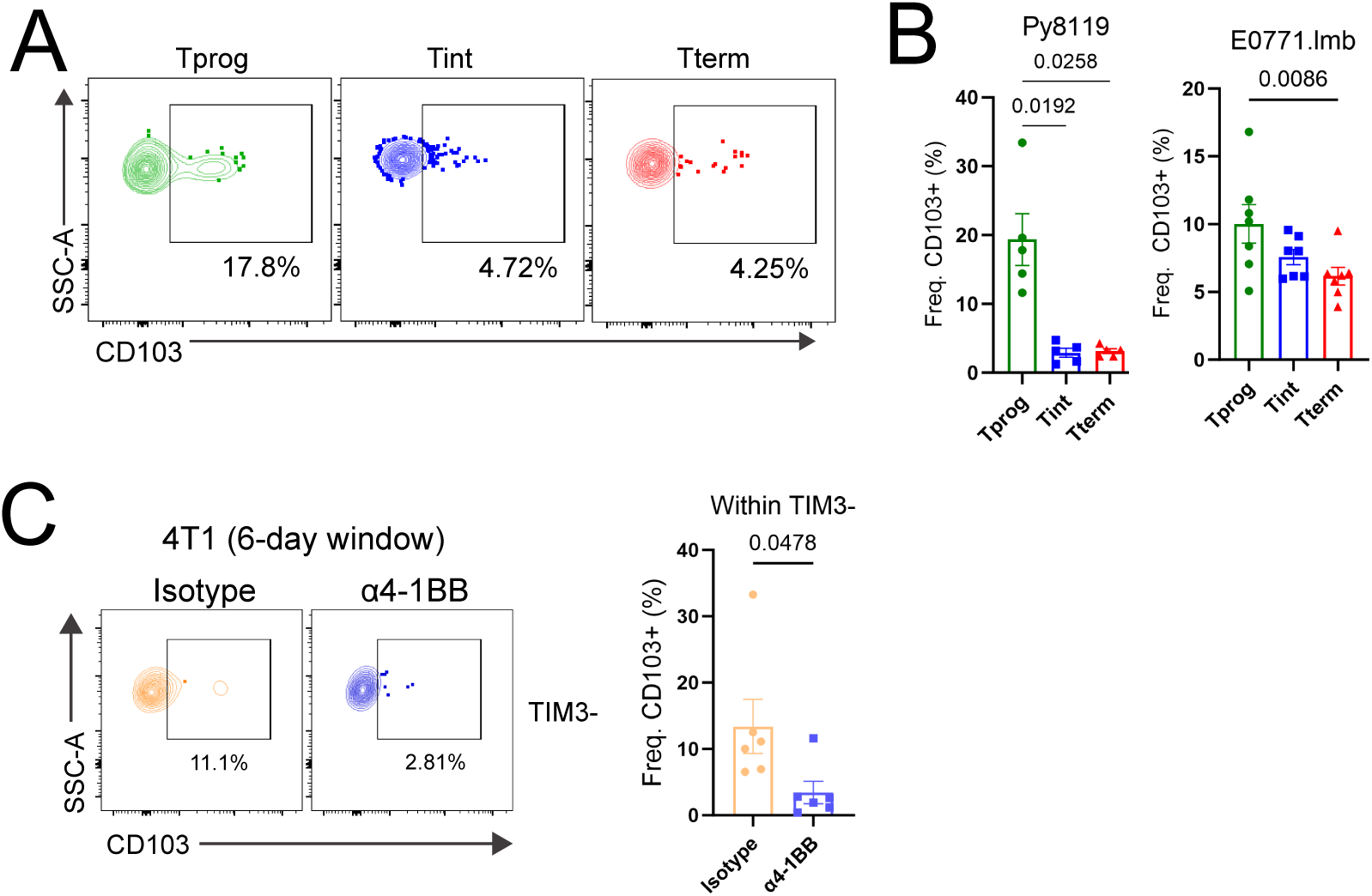
Anti-4-1BB drives reduction of resident marker CD103 in T_prog_. – related to Figure 4. **(A-B)** CD103 shows the highest baseline expression in T_prog_ over Tint and T_term_, representative flow plots (A), quantification of CD103^+^ frequencies (B). **(C)** Representative flow plot showing frequency of CD103 in TIM3^-^T cells treated with isotype or anti-4-1BB (6-day window) in 4T1 (left), quantification (right). Data are presented as mean ± SEM. *P* values were determined using one-way ANOVA followed by post-hoc Tukey’s multiple comparisons test (B), two-tailed unpaired Student’s *t*-test (C).

**Supplementary Figure 7.**
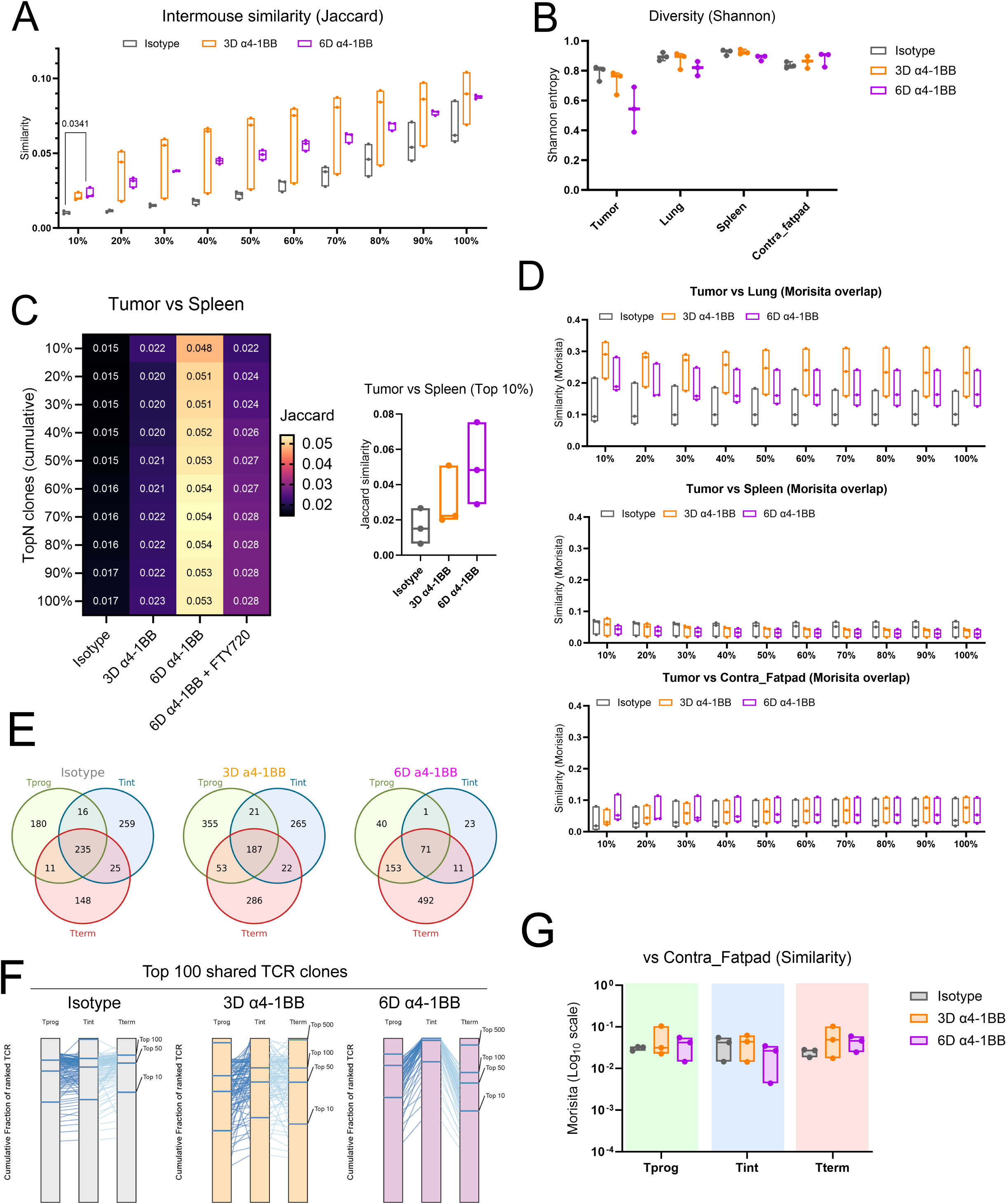
Anti-4-1BB drives expansion of intratumor T cells. – related to Figure 4. **(A)** Jaccard index illustrates 6-day window anti-4-1BB treatment increases TCR similarity between mice. **(B)** Shannon diversity index showing reduction of TCR diversity in tumor under 6-day window anti-4-1BB treatment. **(C)** Heatmap showing T cell similarity between tumor and spleen by Jaccard similarity index. TCRs ranked and cumulatively binned by abundance (left). T cell similarity of top 10% clonotypes between tumor and spleen (right). **(D)** Morisita overlap index shows that anti-4-1BB drives preferential overlap of T cells between primary tumor and lung (top), and not spleen (middle) nor contralateral fatpad (bottom), while considering presence and abundance of unique TCRs. **(E)** Venn diagram depicting overlap of TCR clonotypes between sorted Tprog, Tint and Tterm subsets under all modalities of treatment. **(F)** TCR repertoire analysis shows dynamic changes in clonotype overlap between sorted T cells following anti-4-1BB treatment. **(G)** Morisita overlap index comparing sorted cells and contralateral fatpad harvested from mice under treatments with isotype, 3D α4-1BB or 6D α4-1BB. Data are presented as mean ± SEM. *P* values were determined using Kruskal-Wallis test followed by post-hoc Dunn’s multiple comparisons test (A-D, G).

**Supplementary Figure 8.**
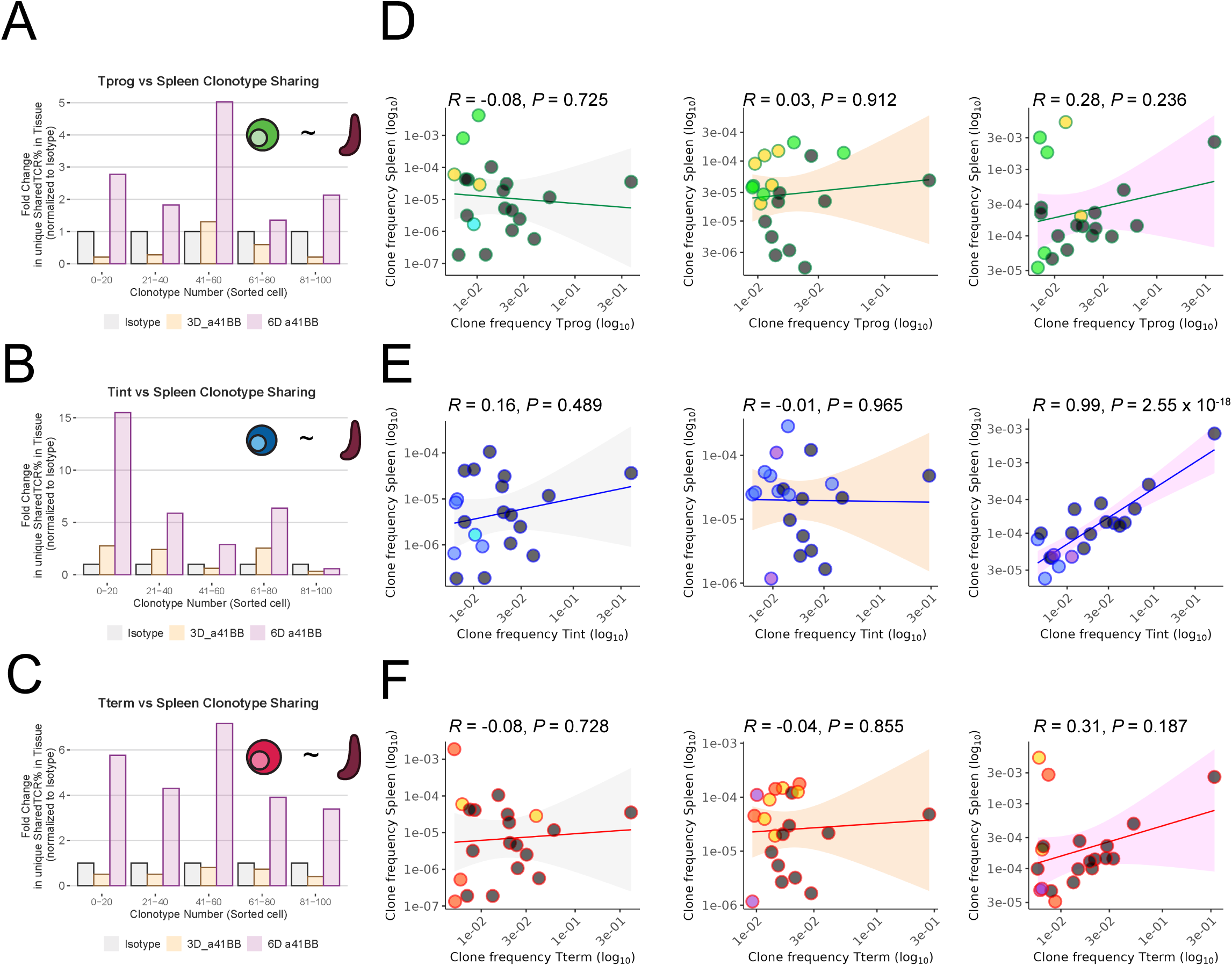
Significant correlation between expanded clonotype in T_int_ and distant tissue related to Figure 5. **(A-C)** Barplot depicting fold change in TCR frequency in the spleen following treatment with either isotype (A), 3D anti-4-1BB (B), or 6D anti-4-1BB (C). Frequencies are normalized isotype frequencies in respective bins. TCRs are first identified from sorted cells, ranked and binned by abundance within the sorted cell groups. **(D-F)** Scatter plots comparing the frequencies of clonotypes in Tprog (D), Tint (E), Tterm (F) to respective frequencies in spleen under Isotype (left), 3D anti-4-1BB (middle) or 6D anti-4-1BB (right) treatment. Correlation coefficient calculated using *cor.test* function. Pearson’s *R* and *P* value was determined using a two-sided t distribution with n − 2 degrees of freedom. Shaded area represents 95% confidence interval of linear model as determined by *geom_smooth* function (grey = Isotype; orange = 3D anti-4-1BB; pink = 6D anti-4-1BB.

**Supplementary Figure 9.**
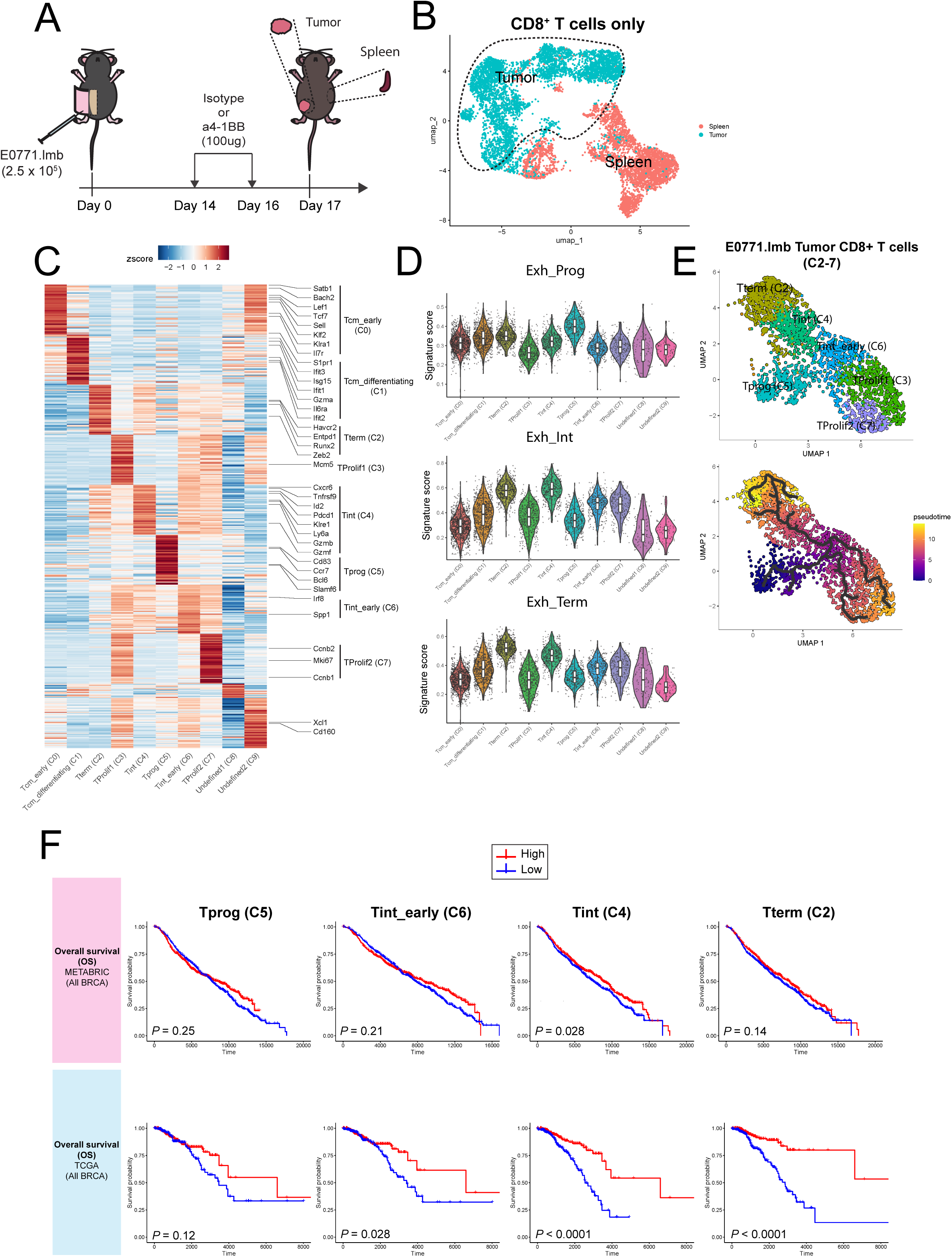
E0771.lmb scRNA-seq and prognostic signatures for overall survival – related to Figure 6. **(A)** Experimental design of E0771.lmb scRNA-seq treated with isotype or anti-4-1BB (3-day window). **(B)** UMAP of total CD8^+^ T cells from both E0771.lmb tumor and spleen. Clusters derived from tumors only (traced by dashed lines) were taken for subsequent CD8^+^ analysis (in Main Fig. 5A). **(C)** Heatmap showing z-score of top 50 variable genes of each cluster, ordered by *p*-value. **(D)** Violin plot showing gene signature of Exh_Prog, Exh_Int and Exh_Term from Giles *et al*. on E0771.lmb CD8^+^ T cell clusters from this study. **(E)** UMAP (top) of CD8^+^ T cells only from E0771.lmb tumors (C2-7) with accompanying pseudotime trajectory (bottom) showing progression of CD8^+^ T cell clusters in development. T_prog_ (C5) was used as the starting point in trajectory analyses. A major branch point was observed in Tint early (C6), with subsequent trajectories proceeding towards Tint (C4) or TProlif1 (C3) and TProlif2 (C7). **(F)** Total BRCA Overall-free survival from METABRIC study (top) and TCGA (bottom) after separating high (3^rd^ quantile) and low (1^st^ quantile) gene signature expression groups. *P* values were determined using log-rank (Mantel-Cox) test (F).

**Supplementary Figure 10.**
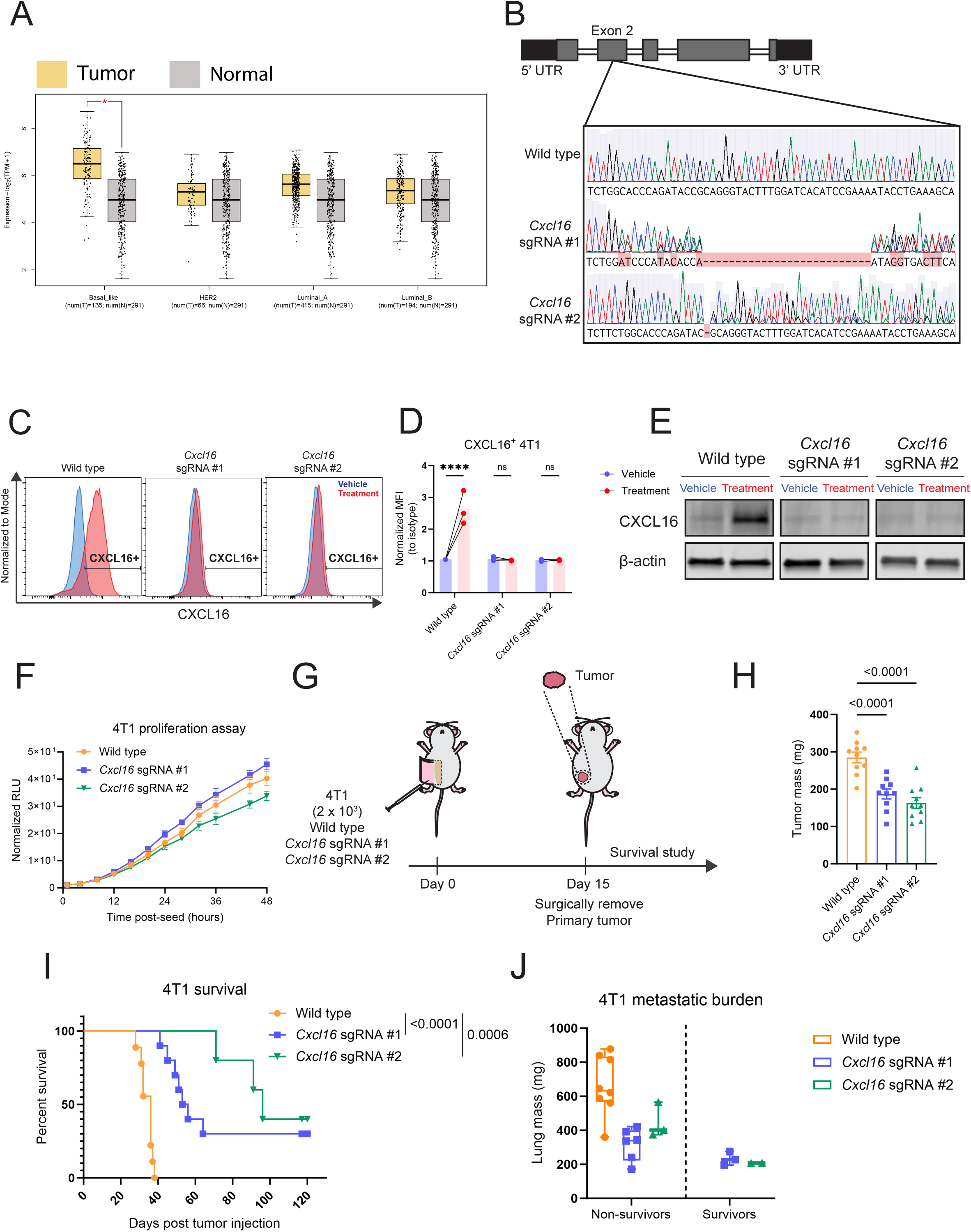
Generation of *Cxcl16KO* tumor model – related to Figure 6. **(A)** GEPIA2 analysis comparing expression of CXCL16 across human breast cancer subtype from TCGA. **(B)** Sanger sequencing results showing successful knockout of *Cxcl16* using gRNA targeting Exon 2. **(C-E)** Protein level verification of knockout using flow cytometry (C and D) and western blot (E). Treatment = 20ng/ml IFNγ + 20ng/ml TNFα + 1µM GI254023X (ADAM10 inhibitor). **(F)** Proliferation assay using Promega Realtime-Flo MT viability assay over a period of 48 hours (data pooled from 3 independent experiments). **(G-J)** *Cxcl16* knockout reduces lung metastatic burden. **(G)** Experimental schematic. **(H)** Primary tumor mass at time of surgical resection. **(I)** Kaplan-Meier analysis. **(J)** Lung metastatic burden (mass) at endpoints. (in H, n = 10 per group; in I and J, wild-type, n = 8-9, *Cxcl16* sgRNA #1, n = 9, *Cxcl16* sgRNA #2, n = 5). *P* values were determined using two-tailed paired Student’s *t*-test (D), one-way ANOVA followed by post-hoc Dunnett’s multiple comparisons test (H), log-rank (Mantel-Cox) test (I).

**Supplementary Figure 11.**
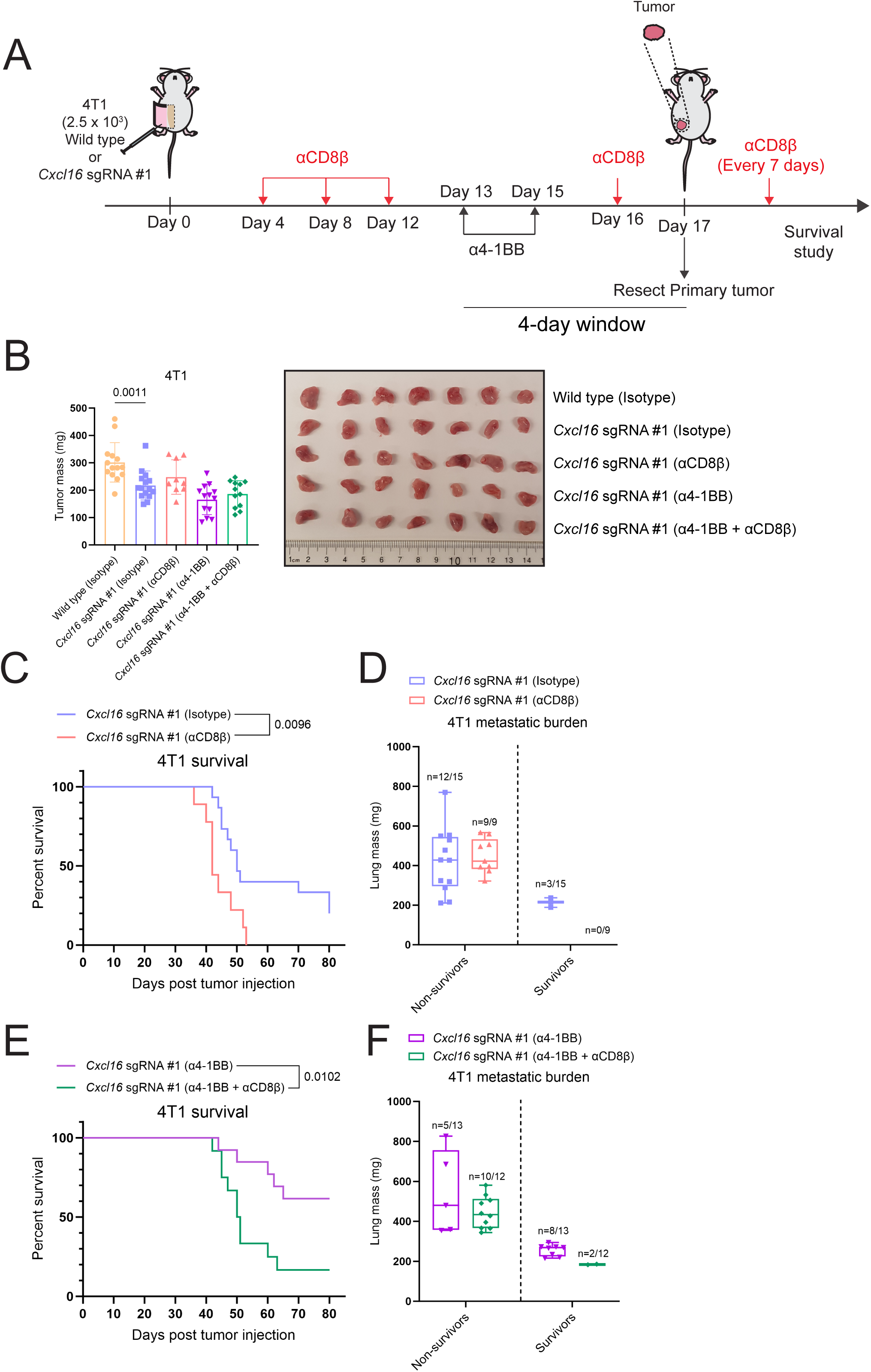
Anti–4-1BB synergizes with loss of *Cxcl16* to mediate survival in a CD8^+^ T cell-dependent manner – related to Figure 6. **(A)** Experimental schematic. **(B)** Primary tumor mass at time of surgical resection (left) and representation image (right). **(C)** Kaplan-Meier analysis of 4T1 *Cxcl16* sgRNA #1 treated with isotype or anti-CD8β. Tumors were surgically resected on day 17 post-tumor inoculation and mice underwent survival analysis. **(D)** Lung metastatic burden (mass) at endpoints for mice in C. **(E)** Kaplan-Meier analysis of 4T1 *Cxcl16* sgRNA #1 treated with anti-4-1BB in combination with or without anti-CD8β. Tumors were surgically resected on day 17 post-tumor inoculation and mice underwent survival analysis. **(F)** Lung metastatic burden (mass) at endpoints for mice in E. (wild type (Isotype), n = 14; *Cxcl16* sgRNA #1 (isotype), n = 15; *Cxcl16* sgRNA #1 (αCD8β), n = 9; *Cxcl16* sgRNA #1 (α4-1BB), n = 13; *Cxcl16* sgRNA #1 (α4-1BB + αCD8β), n = 12). Data are presented as mean ± SEM. *P* values were determined using one-way ANOVA followed by followed by post-hoc Šídák’s multiple comparisons test (B), log-rank (Mantel-Cox) test (C, E).

